# Tricked by Edge Cases: Can Current Approaches Lead to Accurate Prediction of T-Cell Specificity with Machine Learning?

**DOI:** 10.1101/2024.10.23.619492

**Authors:** Martin Culka, Jonathan Desponds, Jeanne Cheung, Mayra Cruz Tleugabulova, Shirley Ng Palace, Martine Darwish, Roman A. Smirnov, Evgeniy Tabatsky, Geraldine Strasser, Andrey S. Shaw, Ira Mellman, Andrei Chernyshev, Darya Orlova

## Abstract

The ability to predict T cell receptor (TCR) specificity from sequence could transform immunotherapy, vaccine development, and our understanding of immune recognition. While machine learning approaches have shown promise, progress is limited by the quality of training data and underlying modeling assumptions. Historically, equilibrium binding assays using multimeric pMHCs have been a dominant source of data, but these assays often conflate high-affinity binding with true functional specificity, introducing noise into predictive models. Here, we critically examine two commonly discussed ideas in the field: that TCR specificity prediction can be separated from functional activation modeling, and that unsupervised sequence-based approaches can generalize across diverse antigen contexts. We introduce a cell-based assay for directly quantifying TCR–pMHC binding kinetics using monomeric ligands, while simultaneously assessing early activation via CD3ζ phosphorylation. These kinetic parameters provide a mechanistic basis for specificity that avoids the artifacts of equilibrium-based measurements. We propose a predictive modeling framework that integrates biophysical measurements with machine learning, and outline strategies for generating high-throughput training data to support this approach. Our findings highlight the need for functionally informed, mechanistically grounded models to advance generalizable TCR specificity prediction.

## Introduction

A few decades ago, multimer-binding technology revolutionized immunology [*Davis et al ., 2011*] by enabling researchers to directly identify and quantify subsets of antigen-specific T cells (i.e., T cells that become activated upon binding to their target). Public databases such as IEDB (https://www.iedb.org/) and VDJdb (https://vdjdb.cdr3.net/) are largely composed of data generated during this “multimer-binding era”. While this technology remains invaluable in certain scenarios, recent insights have revealed its limitations. Over time, multimer-binding technology has biased research toward predominantly higher-affinity TCRs that are not necessarily specific. This bias is evident from two key observations: the growing number of empirical studies revealing a disconnect between multimer-binding capacity and T cell activation (Table S1, Figure 1) and the realization that, while we still lack a clear biophysical definition of TCR specificity (i.e., the parameters of TCR-pMHC interaction that determine whether a T cell becomes activated upon binding to its target), it is evident that affinity alone cannot serve as that measure [*Irving et al ., 2011; Sibener et al ., 2018; Zhao et al ., 2022*]. As a result, assays that measure the binding strength of TCR–pMHC interactions at equilibrium, when decoupled from T cell activation, cannot reliably serve as TCR specificity assays. From a machine learning perspective, training data generated from such binding assays are likely to contain both false positives and false negatives (Figure 1), making it challenging to predict TCR specificity without explicit measurements of T cell activation. Until a clearer biophysical definition of TCR specificity is established, it will remain necessary to rely on training data in which binding assays are coupled with T cell functional readouts.

**Figure 1.**
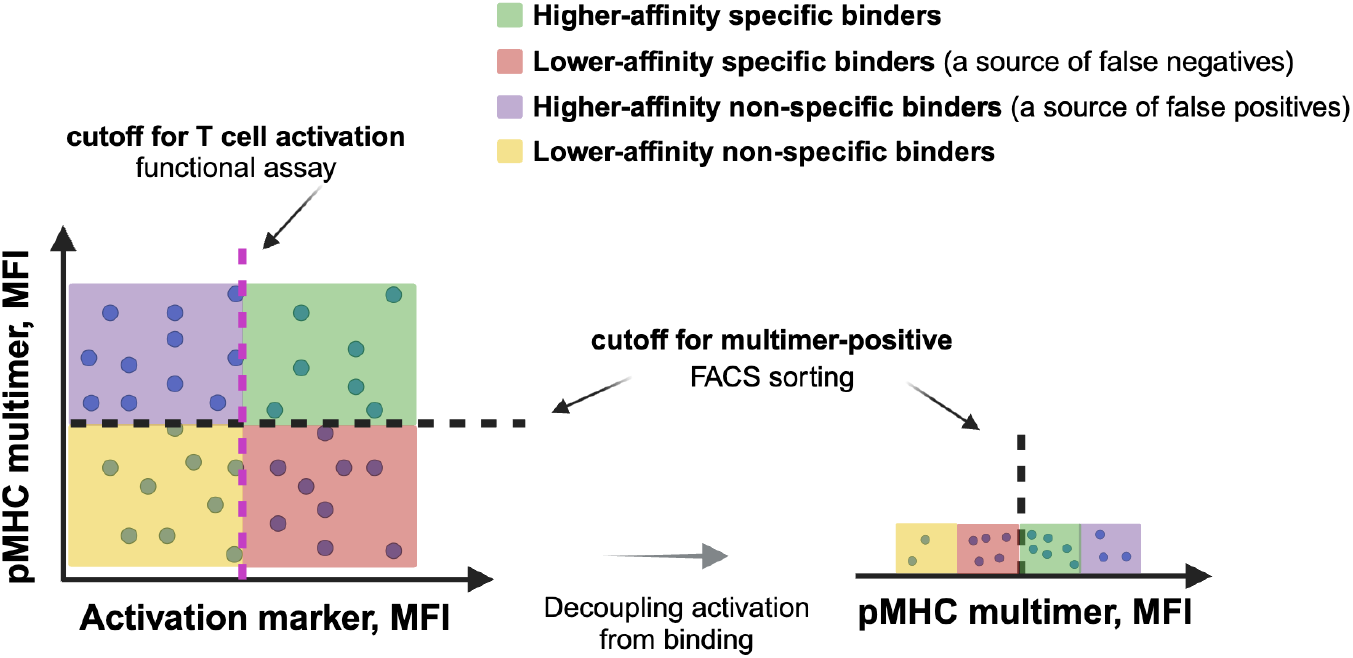
Current multimer-based methods for assessing TCR specificity do not support the separation of specificity and activation prediction into two independent tasks. Omitting functional readouts from multimer-binding assays is effectively equivalent to removing an essential dimension that helps differentiate between specific and non-specific TCR populations. Figure elements created with BioRender.com

That said, while equilibrium pMHC binding, and by extension, affinity, is not a definitive proxy for specificity, pMHC multimers still provide practical value in certain contexts. For instance, using soluble pMHCs in assays where TCRs are presented on cells remains an informative entry point. These assays allow immediate evaluation of co-receptor contributions, such as CD8 and CD4, and enable pairing of binding measurements with assessments of T cell triggering and activation. Nevertheless, existing models in the literature, including kinetic proofreading [*McKeithan, 1995*] and digital signaling [*Zikherman et al ., 2015*], underscore the importance of further investigating the kinetic parameters that govern TCR–pMHC interactions. The kinetic proofreading model proposes that productive T cell activation depends on sustained engagement of the TCR–pMHC complex, during which a cascade of phosphorylation events occurs. This model emphasizes complex half-life (t_1/2_) as a key determinant of signaling. Complementing this, digital signaling models suggest that T cells convert analog inputs (e.g., dwell time, affinity) into binary outputs, reflecting a threshold-based decision-making process. These frameworks are not mutually exclusive and may represent different phases of TCR discrimination.

Nonetheless, direct experimental validation of these models has been hampered by the lack of scalable, quantitative methods for measuring TCR–pMHC binding kinetics in a cellular context. Existing cell-based assays typically rely on multimeric reagents and equilibrium binding measurements, which limit their ability to resolve individual on- and off-rates (*kon* and *koff*) or to capture the dynamics of receptor engagement. Other approaches, such as micropipette adhesion assays [*Huang et al ., 2010*], offer high-resolution insights but are constrained by very low throughput, often allowing measurement from only a handful of cells per experiment. To address this gap, we developed a cell-based assay that quantitatively measures TCR–pMHC interaction kinetics using monomeric pMHC reagents. By leveraging the advantages of fast dissociation rates inherent to monomers, this assay enables precise estimation of *kon* and *koff*, while preserving the physiological context of CD8+ T cells. In parallel, we assess early signaling via CD3ζ phosphorylation to probe the relationship between binding kinetics and functional activation. As we demonstrate here, this framework not only deepens our understanding of TCR–pMHC dynamics but also establishes a foundation for predictive models that integrate mechanistic biophysical parameters, such as kinetic rates, as features for TCR specificity prediction.

In this work, we use the term *“edge cases”* to refer to scenarios in which TCR specificity appears to be adequately captured by simplified descriptors, such as the presence of recognizable sequence motifs or an apparent correlation between binding affinity and functional activation. Although such relationships have been observed and leveraged in prior studies, they arise in restricted regimes and do not represent the general mode of TCR recognition. Outside these special cases, affinity and sequence features alone fail to consistently predict activation, particularly across different antigens, ligands, and experimental conditions. Emphasizing these edge cases can therefore obscure the underlying biophysical determinants of specificity, reinforcing the need for time-resolved, mechanistically grounded measurements of TCR–pMHC interactions that explicitly link binding kinetics to early cellular signaling.

## Results

### Using equilibrium binding as a measure of specificity and as ground truth for predictive modeling

The initial success of equilibrium multimer-binding assays in identifying antigen-specific T cells gave rise to the hypothesis that TCRs with similar sequences, or shared dominant binding motifs [*Glanville et al., 2017*], are likely to recognize the same pMHC. This assumption spurred the development of unsupervised machine learning models that use TCR sequence similarity as a proxy for specificity [*Hudson et al ., 2023; Chronister et al., 2021*]. Recent studies [*Hudson et al., 2024*] that claim these approaches are effective classifiers, despite reporting low discriminatory power, underscore the need for thorough evaluation. Here, we focus on evaluating the conceptual limits of sequence-similarity–based inference under controlled conditions, rather than on optimizing model performance. While our previous study [Culka et al., 2025] is focused on supervised machine learning for TCR–pMHC specificity prediction, the present work complements these findings by examining the limitations of unsupervised approaches and introducing mechanistically grounded measurements to improve predictive modeling.

To assess the validity of this approach, we systematically reanalyzed published clustering data [*Leary et al., 2024*]. (All analyses were performed under HLA-fixed conditions, using the same preprocessing, distance metrics, and clustering procedures described in Materials and Methods.) Our analysis revealed that the ability of these methods to group TCRs by specificity is limited. Only a small subset of TCRs formed pure clusters enriched for a single peptide specificity (Figure 2a), consistent with prior findings reported in Leary et al. [*Leary et al ., 2024*] (Figure 4 and Supplementary Materials). Despite claims of classifier effectiveness [*Hudson et al ., 2024*], the observed clustering patterns demonstrated low discriminatory power, highlighting the need for more robust strategies to predict TCR–pMHC interactions.

**Figure 2.**
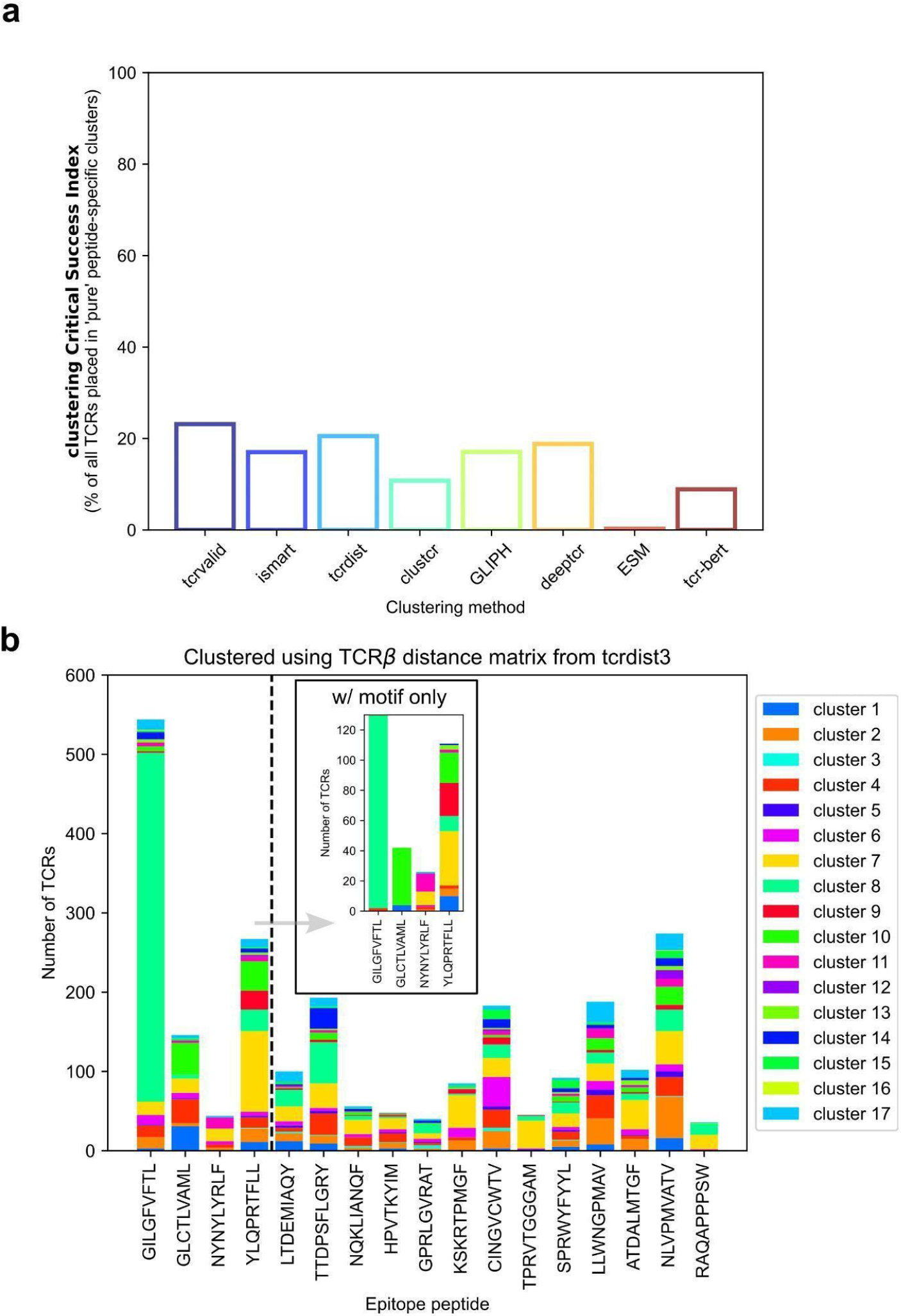
Unsupervised models group TCRs independently of their epitope specificity. **a**. Performance of representative unsupervised machine learning models in sorting TCR sequences into antigen-specific clusters, as reported in Leary et al. [*Leary et al., 2024*]. The plot summarizes clustering outcomes for TCRβ chains using multiple published methods, which together span the most commonly used distance metrics and embedding approaches. Over 70% of datasets result in impure clusters, i.e., clusters that contain mixtures of TCRs specific for different peptides, demonstrating the limited specificity of current unsupervised models. Similar performance trends were observed for clustering based on TCRα or paired TCRαβ sequences (data not shown, see Leary et al. [*Leary et al., 2024*]). **b**. Hierarchical clustering of 17 peptide-specific TCRβ repertoires from the IMMREP_2022 benchmark dataset [*Meysman et al., 2024*], using tcrdist3 distance metric [*Dash et al., 2017; Mayer-Blackwell et al., 2021*]. Each TCR is assigned to one of 17 clusters. The first four repertoires (indicated by the dashed line) correspond to peptides with significant CDR3β sequence motifs, as determined by sequence logo analysis. Despite motif presence, TCRs recognizing the same peptide are distributed across multiple clusters. The inset shows cluster distributions for only those TCRs that contain the dominant sequence motif for each of the four peptides, further illustrating the lack of clear separation based on motif-bearing sequences alone.

To better understand the limitations underlying these observations, we conducted a targeted analysis of motif-based sequence patterns within antigen-specific repertoires. We systematically evaluated peptide-specific TCR repertoires from the IMMREP_2022 benchmark dataset [*Meysman et al ., 2024*] to assess the prevalence and utility of conserved sequence motifs for specificity inference. Conserved binding motifs, as described by Culka et al. [*Culka et al., 2025*], were identified in only 4 out of 17 repertoires (Figure 2b, Figure S1), indicating limited motif occurrence across antigen-specific datasets. Even when present, TCRs containing the dominant motif were distributed across multiple clusters rather than forming a distinct group (Figure 2b, inset), suggesting that motif presence alone does not guarantee cluster homogeneity.

To further evaluate the reliability of sequence similarity as a proxy for antigen specificity, we compared TCR sequence distances using multiple representations, including ESM2 embeddings, plain BLOSUM62 pairwise similarity, and tcrdist3 metrics [*Dash et al., 2017; Mayer-Blackwell et al., 2021*]. These representations span learned embeddings, substitution-matrix similarity, and structure-informed distance metrics, respectively (see Materials and Methods). Across these metrics, we observed that TCRs targeting different peptides frequently exhibited higher sequence similarity to each other than to TCRs recognizing the same antigen (Figure 2b, Figure S1), reinforcing the inconsistency of sequence similarity-based grouping. These findings are consistent with recent observations by Simpson et al. [*Simpson et al., 2024*], where motifs failed to generalize beyond a few specific cases (Figure 6, Figure S11).

Nonetheless, in simpler peptide-specific classification tasks, distance-based clustering approaches performed on par with supervised methods [*Meysman et al., 2024*], highlighting their limited but context-dependent utility. However, their comparable performance in narrow settings does not translate to broader applicability, especially when addressing the full diversity of TCR–pMHC interactions. As a result, the reported performance likely represents an upper bound; introducing MHC/HLA diversity would be expected to further degrade both clustering purity and classifier performance. Introducing variation in MHCs/HLAs within peptide-specific datasets would add further complexity to TCR specificity prediction. Conversely, determining the MHC/HLA type recognized by a TCR may be inherently easier than identifying the specific peptide epitope.

Unsupervised machine learning approaches based on commonly used similarity features, such as binding motifs or readouts from equilibrium multimer-binding assays, are insufficient to fully capture TCR specificity. In our recent study [*Culka et al., 2025*], we also demonstrated the limitations of supervised ML methods that rely on similar data types, including equilibrium multimer-binding assays and, in some cases, functional assay readouts, particularly when applied in a general pan-peptide context. The fact that neither supervised nor unsupervised ML approaches have proven broadly effective in predicting TCR specificity suggests that the underlying data may lack the conceptual resolution required to describe this complex phenomenon. This highlights the need for more advanced strategies, as discussed in the following section.

Together, these analyses indicate that limitations in current ML approaches stem not from model choice, but from the nature of the data used to define specificity, motivating the need for alternative, mechanistically grounded measurements.

### TCR-pMHC binding kinetics as a basis for predicting TCR specificity and enabling modeling

Several decades ago, early efforts to study TCR–pMHC binding kinetics utilized soluble monomeric pMHC molecules with photoaffinity labeling assays [*Luescher et al., 1995*]. Shortly thereafter, flow cytometry–based assays were introduced as a tool to study TCR binding to pMHC [*Altman et al., 1996*]. These newer assays primarily focused on measuring equilibrium binding, where stable interactions are necessary. As a result, monomeric pMHCs were deemed impractical due to their rapid *koff*, which hindered stable binding and accurate affinity measurements. To address this, multimeric pMHC reagents became the standard, offering increased binding stability through avidity effects.

However, for kinetic measurements, a rapid *koff* is advantageous, as it allows direct quantification of both association (*kon*) and dissociation (*koff*) rates - key parameters for characterizing TCR–pMHC interactions. In this context, monomeric pMHCs avoid the avidity-related artifacts introduced by multimeric reagents. Building on this principle, we developed a cell-based assay (Figure 3a) that tracks the binding and unbinding of intact CD8+ T cells to target peptide–MHC monomers using flow cytometry. In parallel, we measured CD3ζ phosphorylation to evaluate the earliest steps of T-cell activation and explore their correlation with TCR occupancy and TCR–pMHC binding kinetics.

**Figure 3.**
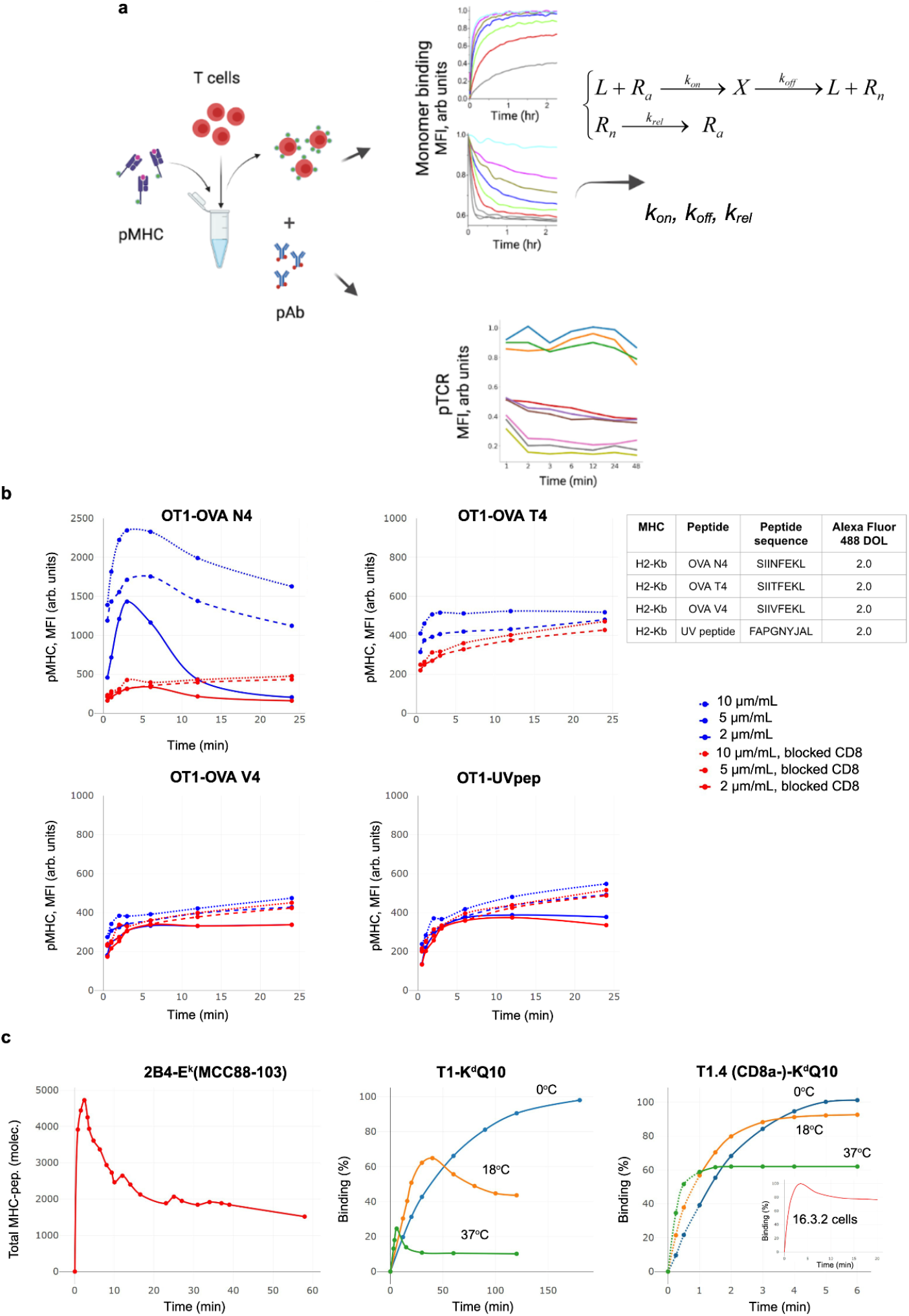
A cell-based monomeric flow cytometry assay enables measurement of TCR–pMHC binding kinetics and recapitulates the previously observed distinctive shape of kinetic curves. **a**. Schematic representation of the assay. Peptide–MHC (pMHC) monomers labeled with Alexa Fluor 488 are incubated with T cells to allow pMHC–TCR binding. Following fixation, cells are stained with phospho-specific monoclonal antibodies (pAb) targeting phosphorylated sites on the ζ domain of the TCR. These pAbs are directly conjugated to APC (indicated by red circles in the schematic). **b**. Representative binding kinetics curves for OT-I T cells interacting with OVA-derived peptides (N4, T4, V4) and a “UV peptide.” Curves are shown with (red) and without (blue) anti-CD8α antibodies, across varying concentrations of pMHC. The degree of labeling (DOL) for each pMHC is reported in the table. **c**. Previously reported TCR–pMHC binding kinetics measured using microscopy [*Grakoui et al., 1999*] (left), photoaffinity labeling assay [*Luescher et al., 1995*] with soluble monomeric pMHC in the presence of CD8 co-receptor (middle), and in the absence of CD8α (right; inset shows T cells expressing CD8α). Figure elements created with BioRender.com

As shown here, this assay successfully recapitulated the previously observed hump-shaped TCR–pMHC–CD8 kinetic curves (Figure 3b). The reappearance of this hump, reported by Luescher et al. [*Luescher et al., 1995*] and others (Figure 3C), and largely overlooked since, is a fortunate observation that serves at least three key purposes: (1) it enables identification of a biokinetic model capable of describing these data by dramatically narrowing the space of possible models; (2) it indicates that a simple reversible ligand–receptor mechanism (i.e., association–dissociation) is insufficient to explain TCR–pMHC–CD8 binding kinetics; and (3) it suggests the presence of a feedback loop, whereby TCR phosphorylation triggers modifications, such as conformational changes, in the extracellular domain of the TCR that in turn affect its binding capacity to pMHC.

We leveraged the distinctive hump in the binding curve to derive a biokinetic model best suited to describe the observed kinetic patterns. Starting with the classical reversible ligand–receptor model (L + R ⇌ LR), we progressively increased model complexity and fit each version to the experimental kinetic data. The simplest model that provided the best fit (Figure 4a) included not only the association and dissociation steps, but also a transition of the receptor into an “inactive” state (Ra → Rn) following its interaction with pMHC, followed by a “relaxation” step back to the “active” state (Rn → Ra) (see Materials and Methods for a detailed description of the “TCR cycle” model).

We further applied this mechanistic mathematical model to a series of well-characterized OT-I–OVA (N4, T4, V4) peptides and compared it to a non-specific peptide (UVpep; see Figure 4b) binding experiments to systematically estimate the underlying TCR–pMHC binding kinetics parameters, including *kon, koff, krel* (see Data S1). This approach provided quantitative insights into the kinetics of each peptide variant and enabled assessment of the impact of anti-CD8α antibodies on TCR–pMHC–CD8 binding kinetics and the resulting level of TCR phosphorylation (Figure 4b; see Data S2 for pMHC concentration-dependent pTCR signal).

**Figure 4.**
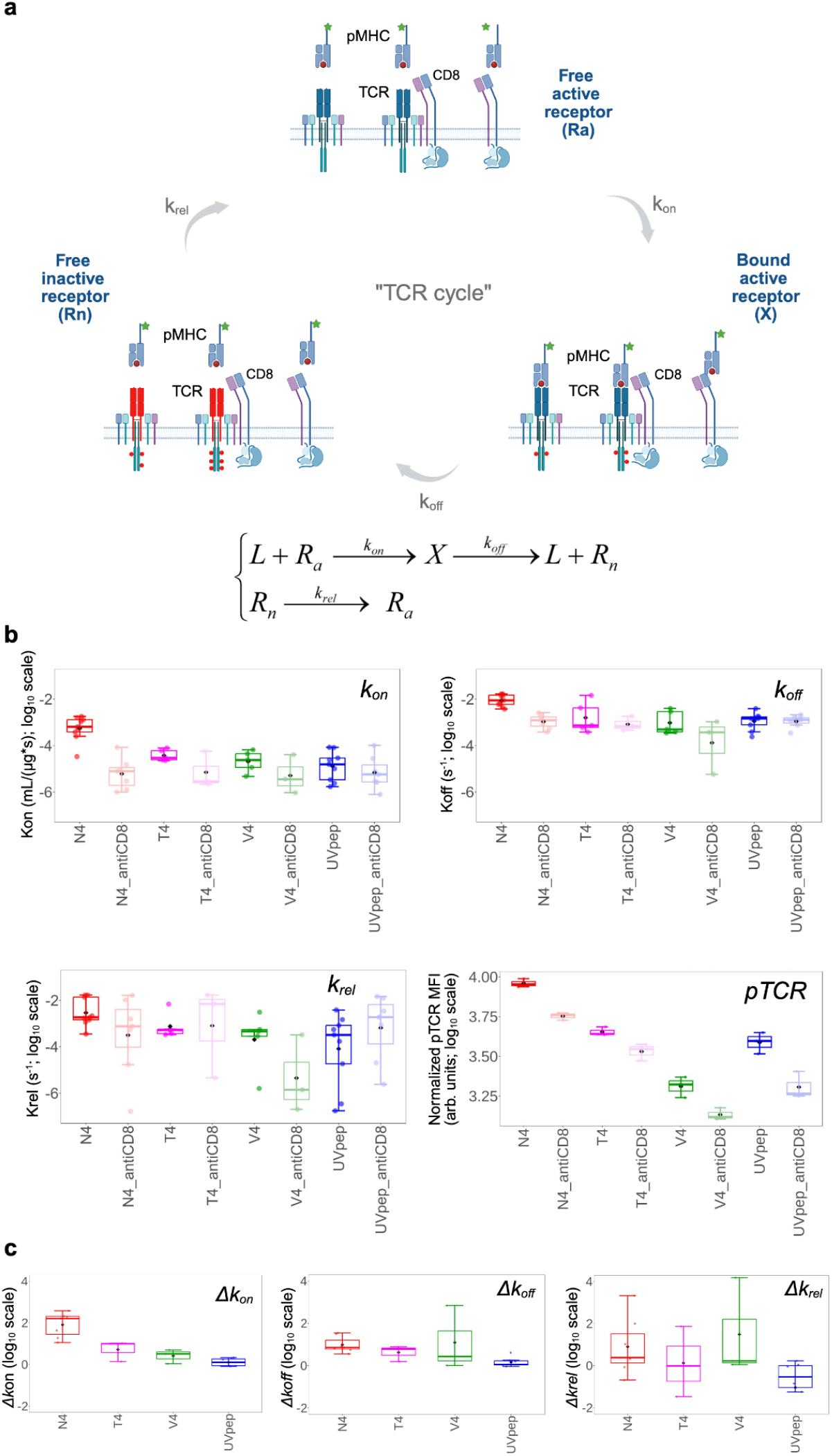
We identified the minimal biokinetics model, referred to as the “TCR cycle”, capable of reproducing the hump-shaped binding kinetics curve. The model (**a**) is consistent with empirical observations (**b**): N4 exhibits higher *kon* than T4, which in turn has higher *kon* than V4. Additionally, the use of anti-CD8α antibodies has the most pronounced effect on N4 (**c**). *k*_*on*_ : Effective association rate of ligand (pMHC) with the active receptor population (R_a_), averaged over TCR-CD8 (higher-affinity), TCR alone (weaker affinity), and CD8 alone (likely negligible). *k*_*off*_ : Effective dissociation rate of ligand-receptor complex (X), reflecting combined off-rates of TCR-pMHC and TCR-CD8-pMHC complexes. Rate of ligand-induced receptor inactivation, captures all routes (e.g., ligand-triggered phosphorylation) by which a bound receptor transitions to a ligand-free inactive state. *k*_*rel*_ : Effective activation rate of receptors, including dephosphorylation, conformational changes enhancing pMHC-binding competency. Rate constant values (*kon, koff*, and *krel*) are presented on a log_10_ scale and reflect data from three or more independent experiments (see Data S1). Phosphorylated TCR (pTCR; CD3ζ phosphorylation) levels are presented as mean fluorescence intensity measured during the early phase of the binding experiment, averaged across three pMHC concentrations, and are used here as an operational readout of early TCR triggering rather than functional avidity. Representative data from one of three independent experiments is shown (see Data S2 for additional datasets). *Δk*_*s*_ : Differences in the corresponding rate constants measured in the absence versus the presence of anti-CD8α antibodies. Error bars represent the standard deviation of values across independent experiments. Figure elements created with BioRender.com

Upon treatment with anti-CD8α antibodies, both *kon* and *koff* values decreased (Figure 4b) relative to untreated conditions. This effect could arise from two factors: first, anti-CD8α antibodies impaired CD8’s ability to participate in TCR–pMHC–CD8 interactions [*Wooldridge et al., 2003*]; second, the antibodies could introduce steric hindrance that affected both binding and unbinding kinetics. The first mechanism is supported by the peptide-dependent patterns in *Δkon* and *Δkoff* (Figure 4c); notably, N4 showed the largest positive *Δkon*, consistent with the empirical observation that N4 was more heavily dependent on CD8 co-receptor engagement than T4 or V4. The second mechanism was corroborated by our steric hindrance test (Figure S2). *kon* decreased due to reduced frequency of productive collisions, as the bulky antibody obstructed proper alignment of TCR, CD8, and pMHC. *koff* also decreased because the steric environment restricted conformational freedom, effectively stabilizing the bound complex and making dissociation less favorable.

We further investigated the role of kinetic parameters in shaping the binding kinetics curve (Figure 5a), emphasizing that the early phase of the binding reaction, occurring over the first few minutes, was the most informative for distinguishing pMHCs based on their specificity for a given TCR (Figure 5b, Figure S3a). This initial phase was primarily governed by the *kon* and *koff* parameters. We also explored the relationship between kinetic parameters and phosphorylated TCR (pTCR) values (Figure 5c), with *koff* showing the strongest correlation with the pTCR signal. This observation is consistent with the TCR cycle biokinetic model, in which *koff* reflects the overall rate of receptor transition to a ligand-free inactive state, including ligand-triggered phosphorylation pathways. Having quantified these kinetic parameters, we next examined how they relate to functional T cell responses.

**Figure 5.**
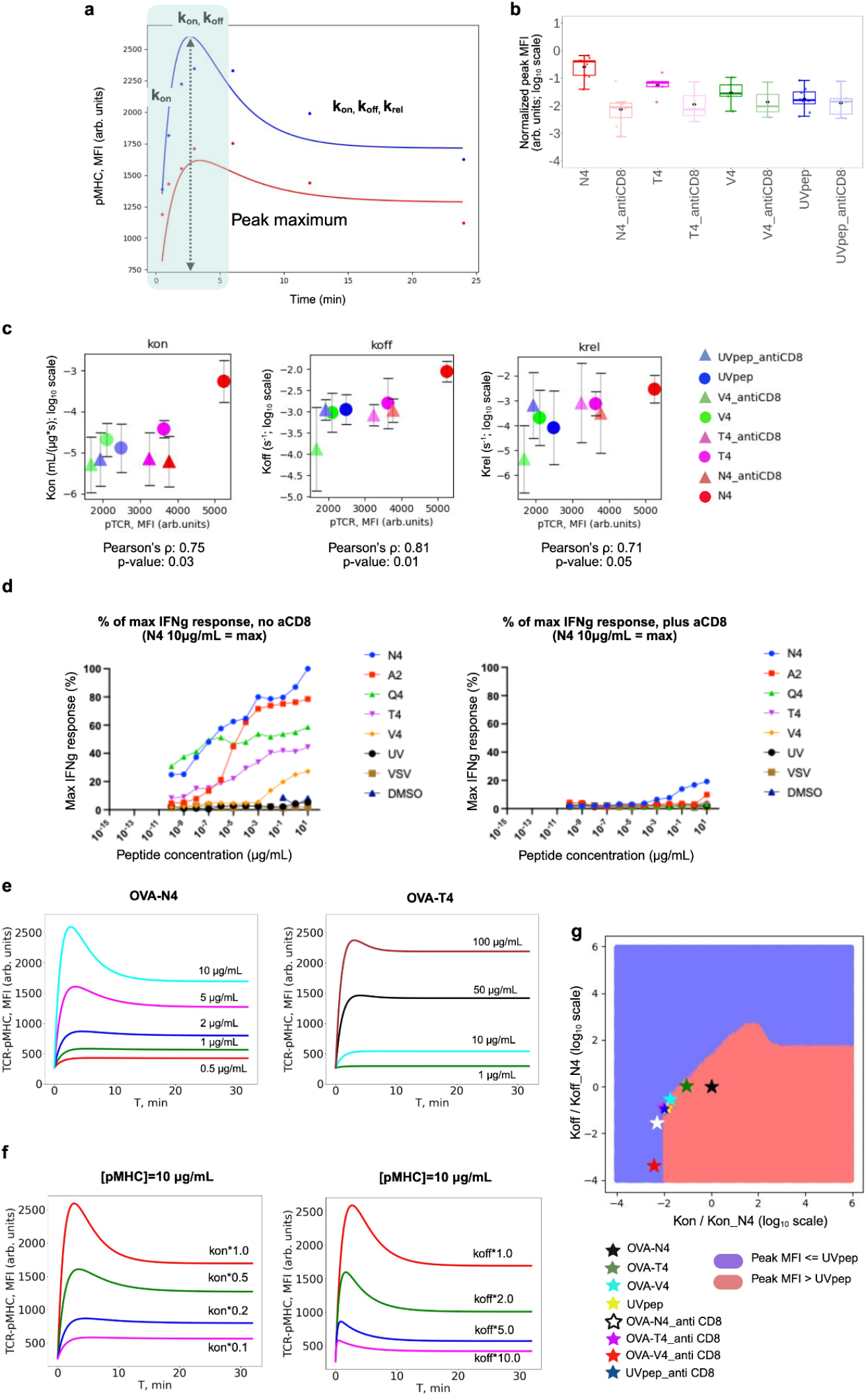
Predictive modeling enables simulation of binding kinetics across varying pMHC concentrations and kinetic parameters. This figure integrates biophysical binding kinetics, early TCR signaling metrics, and downstream functional avidity measurements, with each panel reporting the corresponding operational readout. **a**. Representative binding kinetics data (dots) and corresponding fits using the “TCR cycle” model (solid lines) at two pMHC concentrations (10 µg/mL, blue; 5 µg/mL, red). Key kinetic parameters, *kon, koff*, and *krel* are annotated at the positions on the curve where they exert the greatest influence. For example, *kon* primarily determines the initial slope and intercept of the binding curve at the onset of the TCR–pMHC–CD8 interaction. The dotted line indicates the maximum signal amplitude, while the blue-shaded region highlights the most informative timeframe for distinguishing highly specific peptides from less specific ones. **b**. Maximum (i.e., peak) MFI value observed in the binding kinetics curve during the first 5 minutes of the TCR–pMHC interaction is presented on a log_10_ scale. Values represent averages from three or more independent experiments. To enable comparison across different experimental days, MFI values were normalized using the formula: (peak − B) / A, where A and B represent flow cytometer settings (e.g., voltage), as described in the “TCR cycle” biokinetic model methods section. Error bars represent the standard deviation of values across independent experiments. **c**. Relationship between biophysical kinetic parameters (*kon, koff, krel*) and TCR triggering, quantified by pTCR (CD3ζ phosphorylation) measured during the early binding phase. The Pearson’s ρ and corresponding p-value calculated from mean values are shown in the panel. Pearson correlations calculated from all individual matched data points (not just group means) were as follows: kon Pearson’s ρ=0.65, p-value=0.02, koff Pearson’s ρ=0.49, p-value=0.10, krel Pearson’s ρ= 0.21, p-value=0.53. Error bars represent the standard deviation of values across independent experiments. **d**. Functional avidity assay results for OVA peptide variants measured as cytokine response (IFNγ EC_50_), shown in the absence (left) and presence (right) of anti-CD8α antibodies. Three negative controls (UVpep, DMSO, and VSV) are included. **e**. Binding kinetics curves simulated using the “TCR Cycle” biokinetic model at varying pMHC concentrations, using *kon, koff*, and *krel* parameter values corresponding to N4 (left) and T4 (right). **f**. Binding kinetics simulated at a fixed pMHC concentration for varying *kon* (left) and *koff* (right) values. Here, *kon* *1 refers to the empirically measured *kon* for the OTI–OVA–N4–CD8 interaction, as identified in our manuscript. Accordingly, *kon*\*0.5 represents half that value, and so on. The same convention applies to *koff* (right). **g**. Kinetic parameter space (*kon, koff*) showing regions that produce peak MFI values equal to or lower than the OT-I–UVpep peak (violet area) and those producing higher peaks (red area). Peak MFI is used here as an early signaling metric. Parameter values are shown relative to N4 on a log_10_ scale. Simulations were performed at a pMHC concentration of 10 µg/mL. Positions of N4, T4, V4, and UVpep are indicated by colored stars.

The *Δkoff*-based ranking of peptides (Figure 4c) aligns well with the functional avidity results (Figure 5d). In addition to reflecting the extent of CD8 involvement in peptide recognition, it suggests that if binding kinetics were measured between OT-I TCRs and the OVA-A2 or -Q4 peptides, the resulting curves would likely still exhibit humps, with peak MFI values falling between those observed for N4 and T4. More broadly, having an analytical solution enables predictive modeling of binding curve behavior in response to changes in pMHC concentration (Figure 5e), and even allows simulation of how the curves would shift with changes in kinetic parameters (Figure 5f, Figure S3b-d). This framework opens the opportunity to computationally explore which combinations of biokinetic parameters yield binding curves that resemble those of highly specific TCR–pMHC interactions, such as OTI–OVA–N4 (Figure 5g, Figure S3b-d), and to identify the optimal pMHC concentrations for TCR–pMHC pairs with such kinetic profiles.

Consistent with these insights, early binding features (0–5 min) more accurately predict TCR functional avidity than late-time binding measurements (20–30 min), highlighting the importance of initial association kinetics for discriminating peptide variants (Figure S4). Correlations between *kon, koff*, and *krel* with functional avidity further support the mechanistic relevance of these kinetic parameters in shaping T cell responses.

Although *kon* shows the strongest individual correlation with functional avidity (Figure S4), it is clear that the observed relationship arises from a combination of kinetic parameters, including koff and krel, underscoring that T cell responsiveness is shaped by the integrated dynamics of the TCR–pMHC interaction rather than any single rate constant.

To further assess the generality of the TCR cycle model, we applied it to previously published murine and human TCR–pMHC systems (Figure 6). Importantly, these datasets were generated by independent laboratories and were not used in any way to develop or tune the TCR cycle model, providing an unbiased external validation. Kinetic traces were extracted from these studies using a standardized digitization workflow, and the model was fitted to association time courses. Across diverse systems and experimental conditions, the TCR cycle model accurately recapitulated the temperature dynamics of binding, including effects of co-receptor engagement and peptide-specific variations. In particular, inferred koff values from the model strongly correlated with dissociation-only measurements, supporting the robustness of the model for capturing intrinsic receptor–ligand kinetics without functional readouts.

To test whether the additional ‘competence cycling’ step (*krel*) is mechanistically required, we compared the TCR cycle model to a simpler two-state reversible model. Across ligands and experimental conditions, the full TCR cycle model consistently provided superior fits (higher R^2^) and lower AIC values, indicating that the observed binding kinetics, particularly hump-shaped curves, cannot be explained by a simple reversible interaction alone (Figure S5a–b). Examination of the fitted parameter correlations further confirms that *kon, koff*, and *krel* are identifiable within the experimental time windows and concentration ranges used (Figure S5c), supporting the mechanistic relevance of competence cycling in shaping TCR–pMHC dynamics. Notably, one of the systems analyzed in Figure 6 exhibits non-monotonic (‘hump-shaped’) association kinetics even when peptides are presented on antigen-presenting cells (Figure 6b), supporting the physiological relevance of the kinetic behavior captured by the TCR cycle model.

**Figure 6.**
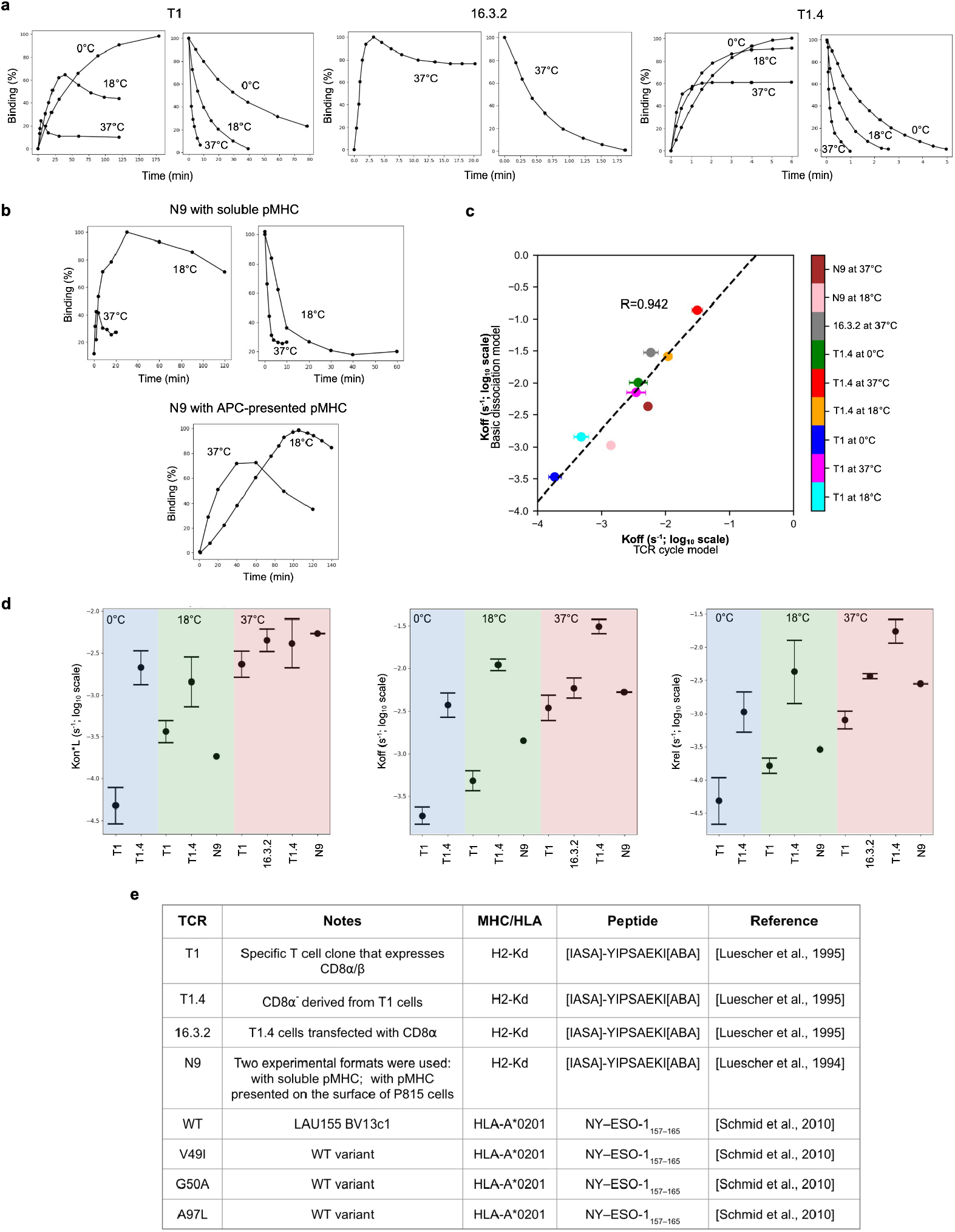
Validation of the TCR cycle model using previously published TCR–pMHC systems. All panels report biophysical binding kinetics, with no functional or signaling readouts included in this figure. **a–b**. Representative association and dissociation kinetic curves for murine TCR–pMHC systems investigated in this study, averaged over three experimental replicates. The datasets were drawn from previously published studies and include measurements performed under different temperature regimes, as indicated. **c**. Correlation between the dissociation rate constants (*koff*) inferred from fitting the association time courses using the TCR cycle model and the *koff* values obtained by fitting dissociation-only kinetic measurements with a standard single-exponential model. Strong agreement is observed (Pearson R > 0.94). Error bars represent the standard errors of the fitted parameters, derived from the Hessian matrix of the fit. **d**. Fits of the TCR cycle model to the representative association kinetic curves shown in panel a, demonstrating the model’s ability to capture the full temperature dynamics of TCR–pMHC binding, as well as the biological effect of mutating the T1 TCR to disrupt CD8α engagement (see panel e for details). Error bars represent the standard errors of the fitted parameters, derived from the Hessian matrix of the fit. **e**. Summary of the murine and human TCR–pMHC (HLA) systems extracted from the published literature and analyzed in this manuscript, including receptor–ligand identities and experimental conditions. As raw numerical data were not publicly available, kinetic traces in panels a–b were digitized from the original publications listed in panel e using custom, open-source tools developed by the authors (see Code availability), ensuring consistent and reproducible extraction across studies.

## Discussion

Leveraging machine learning for TCR specificity prediction is an ambitious and promising goal. However, success critically depends on asking the right biological questions, framing the problem in a way that machine learning can meaningfully address, and generating training datasets that are well-aligned with that formulation. As we have shown in our prior work [*Culka et al., 2025*] and in this study, currently available datasets, primarily derived from equilibrium TCR-pMHC binding assays, sometimes supplemented with T cell activation readouts, remain insufficient for enabling accurate TCR specificity prediction in the general case, using either supervised or unsupervised machine learning approaches. This insufficiency stems not only from limited dataset size (i.e., the number of characterized TCR-pMHC pairs) but also from fundamental limitations in the type of data being collected, and perhaps even in the questions being asked.

Using the well-characterized OT-I/OVA system, we demonstrated that TCR–pMHC interactions are fundamentally kinetic in nature, shaped by a complex interplay between kinetic parameters, receptor and ligand concentrations, and intracellular feedback mechanisms, which collectively manifest as an effective adaptive behavior of the TCR at the systems level. Therefore, rather than relying solely on affinity or avidity measurements at equilibrium, it may be more informative to assess TCR specificity based on early binding kinetics. Importantly, competence cycling should not be interpreted as evidence for a specific extracellular conformational switch. Rather, it represents a minimal systems-level description of feedback and adaptation required to explain the observed kinetics, which could arise from multiple intracellular processes, including phosphorylation-dependent signaling, kinase–phosphatase balance, or receptor–co-receptor coupling. Accordingly, the OT-I–OVA system is used here as a well-controlled validation platform, while the mechanistic framework itself is intended to be broadly applicable to TCR–pMHC interactions across receptors, peptides, and species. Practically, for empirical selection of specific TCRs, this suggests that direct measurement of TCR–pMHC binding kinetics may not be necessary. Instead, existing tetramer staining protocols could be adapted: for example, by reducing the incubation time to 3–5 minutes and using lower concentrations of tetramers (or monomers), rather than the conventional >20–30 minute incubation. In general, such cell-based assays provide a strong starting point for TCR selection and training data generation, as they inherently account for co-receptor involvement and the various feedback mechanisms present in living cells that influence how TCRs discriminate between pMHCs. Consistent with this feedback-based interpretation, prior studies [*Wooldridge et al., 2009; Lissina et al., 2009*] using Src-family kinase inhibition (e.g., dasatinib) report selective stabilization of surface TCR–pMHC interactions, an observation that aligns with the competence cycling framework explored here.

For example, OVA peptide variants (e.g., N4, Q4, T4, V4) differ in their MHC binding and recognition strength by OT-I TCR. Both the functional avidity assays and binding kinetics parameters independently suggest that, at least in the OT1–OVA system, peptide discrimination primarily relies on the CD8 co-receptor. The co-engagement of CD8 significantly enhances the OT1–OVA interaction, both increasing the *kon* and facilitating signal transduction. CD8 plays a crucial role in amplifying the differences between the OVA peptides (e.g., N4, T4, V4) making it essential for fine-tuned discrimination. Does the strong dependence on co-receptor involvement for peptide discrimination make predicting TCR specificity from sequence a hopeless task? Not necessarily. As we have shown here, the only differences between the TCR–pMHC pairs were single amino acid substitutions among N4, T4, and V4. This clearly suggests that, with sufficiently large and well-structured training data, it should be possible to detect patterns of amino acid combinations on both the TCR and pMHC sides that govern CD8 involvement. What is less straightforward to address with machine learning is the fact that, by default, there is no simple mapping from amino acid sequences of the TCR and pMHC to definitive T cell specificity or functional response. Whether a T cell becomes activated in response to a given TCR–pMHC interaction depends significantly on the interplay between TCR and pMHC concentrations. So, how can machine learning help in this context? One promising approach is to integrate machine learning with predictive mechanistic modeling. As we demonstrated, having an analytical solution enables the simulation of binding curve behavior in response to changes in pMHC concentration. However, these predictions rely on empirically measured biokinetic parameters such as *kon* and *koff*. Developing machine learning methods that can approximate *kon* and *koff* from the amino acid sequences of TCR and pMHC could be highly valuable, providing critical inputs for downstream predictive modeling. An immediate question that arises is how to generate sufficient training data to develop an ML model capable of approximating *kon* and *koff* from TCR–pMHC sequences. One potential scalable approach is to apply the binding kinetics measurement framework we introduce here to a diverse repertoire of TCRs, a library of pMHCs, or even a library-on-library setup. This could be achieved by barcoding different incubation time points, representing distinct stages of the TCR–pMHC interaction, with tags detectable by flow cytometry or sequencing. Subsequent sequencing of the TCRs would enable reconstruction of their binding kinetics profiles across time points, allowing *kon* and *koff* values to be extracted by fitting the relevant biokinetic model to these profiles.

From the TCR–pMHC interaction perspective, predicting kinetic parameters (*kon, koff*) from sequence is a more tractable and generalizable modeling approach than predicting functional avidity directly. Kinetic measurements are grounded in biophysics and available across diverse receptor–ligand systems, enabling larger, more consistent training datasets. In contrast, functional avidity data are difficult to scale and standardize across contexts (multiple TCRs, antigens, and cellular contexts). Focusing on kinetics also enables rational receptor design with tunable binding dynamics, providing both predictive insight and translational utility. In our work, we outline a potential path for linking these kinetic parameters to downstream functional outcomes such as functional avidity, providing a conceptual and computational bridge between biophysical binding data and biological function. The large-scale *kon* and *koff* measurements enabled by our multiplexed platform will provide the supervised training labels necessary to develop machine-learning models capable of predicting TCR–pHLA kinetic parameters directly from sequence information. In this framework, experimentally derived *kon*/*koff* values will serve as regression targets, while model inputs will consist of TCR and pHLA sequence embeddings together with sequence-to-structure–derived interface features. This strategy builds on recent advances in computational and machine-learning approaches for predicting biomolecular binding kinetics from structural and biophysical representations, including graph-based and hybrid physics-informed models (reviewed in [*Wang et al., 2023*]). Once trained on our systematically generated kinetic dataset, such models will enable scalable inference of TCR–pHLA binding kinetics across diverse sequence repertoires and will be rigorously validated using held-out experimental measurements.

Looking ahead, a multiplexed library-on-library approach (Supplementary Materials, Figure S6) could enable pooled measurement of hundreds of TCRs against hundreds of pHLA ligands, with time-resolved barcoding allowing reconstruction of association and dissociation kinetics and estimation of *kon, koff*, and *krel* at much larger scale.

More broadly, the cell-based TCR–pMHC kinetics approach we present here opens new opportunities to explore fundamental questions about T cell activation mechanisms. For example, Wooldridge et al. [*Wooldridge et al., 2009*] reproduced an intriguing empirical observation initially reported by Lissina et al. [*Lissina et al., 2009*]: incubation with the protein tyrosine kinase inhibitor dasatinib paradoxically enhances the stability of surface TCR–pMHC interactions, significantly improving staining of cognate T cells with pMHCI tetramers. Dasatinib blocks phosphorylation of the CD3ζ chain and ZAP70, which are early key steps required for T cell activation. These observations align with our findings and the TCR cycle model, which posits that phosphorylation of intracellular TCR domains acts as a feedback mechanism to promote ligand dissociation, thereby increasing *koff* as defined in the model. Within the TCR cycle model, kinase inhibition is predicted to reduce the effective activation rate into the inactive (post-engagement) state, while phosphatase inhibition would be expected to reduce *krel*, prolonging the competent state. These qualitative predictions provide a concrete experimental route for distinguishing among molecular implementations of the feedback captured by the model. It would be valuable to revisit these studies in depth and incorporate protein tyrosine kinase inhibition into future experiments within the TCR cycle model framework.

## Materials and methods

To clarify terminology used throughout this study, we define the following operational metrics: affinity refers to the strength of the TCR–pMHC interaction, typically quantified by the dissociation constant (Kd) or the association/dissociation rates (*kon*/*koff*); specificity refers to the ability of a TCR to selectively recognize its cognate pMHC over unrelated pMHCs; and functional avidity captures the overall potency of a T cell’s functional response to a given pMHC, operationally measured here by readouts such as cytokine EC_50_ (e.g., IFNγ) or EC_50_ for cell-mediated lysis. These definitions will be used consistently throughout the Results to link measured biophysical parameters to functional outcomes.

### Previously published data overview

For the assessment of the peptide-specific predictive power of various clustering methods (Figure 2a), we used data from the Source Data table in Leary et al [*Leary et al., 2024*].

For the assessment of peptide specificity assignment using agglomerative clustering (Figure 2b), based on various distance metrics, we used the IMMREP_2022 benchmark data set (Meysman et al. [*Meysman et al., 2024*], available at https://github.com/viragbioinfo/IMMREP_2022_TCRSpecificity), which contains 17 peptide-specific data buckets. For our clustering analysis, we used concatenated training set data, organized epitope-wise.

### Analysis of previously published data

For Figure 2a, we plotted a subset of an already published analysis (Leary et al. [*Leary et al., 2024*]). Specifically, the data can be found in *fig4cd_sup_fig_1to7* tab of the Source Data table. We selected a subset of data points with minimal cluster size 3, without a spike of irrelevant data (*spike_x* = 0), *tcrvalid* reference labels, and clustering based on TCR*β* (*TRB*). For the methods where it is defined (all except GLIPH and clustcr), we selected data points for the distance parameter *ε*=0.5 (or the closest value available). For each method, we plotted the value in the c-CSI column (clustering Critical Success Index).

For Figure 2b and Figure S1, we clustered aggregated peptide-specific TCRs using agglomerative (hierarchical) clustering with the *scikit-learn* [*Pedregosa et al ., 2011*] Python library. We used distance matrices based on the tcrdist3 [*Dash et al ., 2017; Mayer-Blackwell et al ., 2021*] package (TCR*β* only), Euclidean distance in the latent space of the ESM2 language model [*Lin et al ., 2023*] (*esm2_t36_3B_UR50D* variant, using mean representations of CDR3*β* with 2560 dimensions) and negative BLOSUM62 pairwise sequence similarity (using *pairwise2*.*align*.*globaldx* in BioPython package [*Cock et al ., 2009*]). We then divided the data by epitope and plotted the portions corresponding to each cluster within each epitope-specific data bucket. The subset of motif-containing peptides was selected based on sequence logos (created using logomaker [*Tareen et al ., 2020*]) of CDR3*β* sequences, aligned based in IMGT numbering using ANARCI [*Dunbar et al ., 2016*].

### pMHC reagent generation and characterization

#### UV-MHCI generation and fluorophore labeling

Recombinant H2-Kb heavy chain and B2M were refolded in the presence of a high affinity peptide with the sequence FAPGNYJAL, with residue “J” denoting a non-natural UV cleavable amino acid as previously described [*Bailey, 2013*]. The resultant UV-MHCI was incubated with a 2-20X molar excess of Alexa Fluor 488 NHS ester (ThermoFisher, Cat # A20000) in phosphate buffered saline pH 7.4 for 2 hours at room temperature. Excess, unconjugated fluorophore was removed by dialysis. The sample was loaded into a 10K molecular weight cutoff dialysis cassette (Slide-A-Lyzer 10K MWCO cassette, Thermo Fisher) and placed in 25 mM TRIS pH 8.0, 150mM NaCl, 4 mM EDA at a ratio of 1:2000, sample:dialysate. Sample and dialysate were incubated at 4°C for 8 hours while continually mixing via a magnetic stir plate. Dialysate was then discarded and replaced for an additional 8 hours incubation at 4°C, while stirring. After the second round of dialysis, the sample was recovered, and protein concentration was determined using a UV-Vis spectrophotometer, and was corrected for the contribution of the fluorophore to the absorbance at 280 nm.

#### Determining degree of MHCI fluorophore labeling

The degree of fluorophore labeling (DOL) was determined by reversed phase liquid chromatography mass spectrometry (RP LC-MS). 2-3 ug of the MHC-fluorophore conjugation reaction were injected on an Agilent 1290 Infinity series HPLC in line with an Agilent 6230 time-of-flight electrospray ionization mass spectrometer. The reversed phase column (Agilent PLRP-S 1000 Å, 8um, 50 x 2.1 mm) was exposed to a gradient of 25-45% mobile phase B in 5 min at 0.50 ml/min with the column heated to 80°C. Mobile phase A was 0.05% TFA and mobile phase B was 0.05% TFA in acetonitrile. The column eluent was sent to the TOF-MS for mass spectrometry data acquisition. The degree of fluorophore conjugation was determined by using the deconvoluted mass spectra of the peaks corresponding to B2M and H2-Kb heavy chain. The average number of fluorophores per pMHCI was calculated by using the relative abundance of each mass corresponding to a different number of fluorophore additions to each protein species and determining a weighted average for the complex.

#### UV-mediated peptide exchange

OVA synthetic peptides (Anaspec, Elim Bio) were solubilized in ethylene glycol to a concentration of 20 mg/ml and were added to fluorophore labeled peptide-MHCI at a 25X molar excess. The peptide exchange reaction was performed in 25 mM TRIS pH 8.0, 150 mM NaCl, 4 mM EDTA, and contained 5% ethylene glycol v/v after the addition of peptide. The final concentration of the peptide-MHCI in the exchange reaction was 2.0 mg/ml. The peptide-exchange reaction was then incubated under a UV light set to 365 nm (Analytikjena UVP 3UV Lamp) for 20 minutes. After exposure to UV light, the exchange reaction was allowed to proceed at room temperature for a minimum of 4 hours, or overnight incubation.

#### Determination of peptide binding to MHCI

A 2-dimensional liquid chromatography mass spectrometry (2D LC-MS) method was used to characterize peptide binding to MHCI complexes. Between 2-3 ug of MHCI-peptide mixtures were injected on the instrument and sent to the first dimension column. The first dimension LC method employed an analytical size exclusion column (SEC) (Agilent AdvanceBio SEC 300Å, 2.7um, 4.6 x 15 mm) to separate intact complex from excess peptide run at an isocratic flow of 0.7 ml/min in 25 mM TRIS pH 8.0, 150 mM NaCl for 10 min with signal acquisition at 280 nm. A sampling valve collected the entirety of the complex peak corresponding to monomeric pMHCI that eluted between 1.90 – 2.13 min in a volume of 160 ul and injects it onto the second dimension reversed phase column (Agilent PLRP-S 1000 Å, 8um, 50 x 2.1 mm). The second dimension column was exposed to a gradient of 5-50% mobile phase B in 4.7 min at 0.55 ml/min with the column heated to 80°C. Mobile phase A was 0.05% TFA. Mobile phase B was 0.05% TFA in acetonitrile. The column eluent was sent to an Agilent 6224 TOF LCMS for mass spectrometry data acquisition.

MHC-I complex peak area in the first dimension and mass spec detection of the peptide in the second dimension were used to determine successful peptide binding. Successful binding of a peptide into the complex after cleavage of the conditional ligand during the peptide exchange reaction stabilizes the complex and results in nearly complete recovery of the starting complex measured in the first dimension SEC analysis. The peptide that has exchanged into the complex can then be detected in the second dimension, where the complex was run under denaturing conditions with mass spectral analysis, allowing for direct detection of the peptide of interest. The first dimension SEC method also allowed for the detection of any aggregate species. Aggregation formation during the peptide exchange reaction is most often formed from partially unfolded HLA from a complex that failed to bind a peptide. The pMHCIs used in this study were >98% monomeric and are unlikely to have contained any functional higher order complexes.

### Flow cytometry based association and dissociation kinetics assay (including pTCR)

OTI CD8 T cells were purified by negative selection (Miltenyi Biotec, 130-104-075) from spleens and lymph nodes of OTI mice (Jackson Laboratory or Genentech in-house colony). All animal usage was conducted by following relevant ethical regulations detailed in animal use protocols approved by the Genentech Institutional Animal Care and Use Committee. Cells were stained with live/dead Aqua (Thermo Fisher Scientific, L34966) and TCRb BV711 (BioLegend, clone H57-579) prior to dispensing in a 96 well plate at 3-4x10^5/well. For association assays, titrated amounts of peptide:MHC monomers were added at 4C, and at specified time points, 4% paraformaldehyde fixation buffer was added. For dissociation assays, titrated amounts of peptide:MHC monomers were prebound to cells at 4C for 1 hour, washed, and then fixed at specified time points.

For CD8 blockade conditions, purified anti-CD8 (Thermo Fisher Scientific, clone MA5-17594) was prebound at 10 µg/mL and then at 50 µg/mL in the presence of the peptide:MHC monomers. Association and dissociation assays were run in MACS buffer (1x PBS + 0.5% BSA + 2mM EDTA) at 4C. To detect the level of phosphorylation of the ζ domain of TCR, cells were washed with 1x permeabilization buffer (Thermo Fisher Scientific, 00-8333-56) prior to incubation with pCD247 CD3ζ Tyr142 APC (Thermo Fisher Scientific, 17-2478-42) in 1x permeabilization buffer for 30 minutes at room temperature. Cells were then washed with 1x perm buffer and resuspended in MACS buffer for subsequent flow cytometric analysis. Samples were run on a BD Symphony flow cytometer, followed by data analysis using FlowJo software.

### Association kinetics assay automation

Plate washing and fixation procedures were performed using a fully integrated automation platform developed by HighRes Biosolutions, operated under Cellario scheduling software. The system was configured to execute time-sensitive liquid handling and environmental control steps with high precision and reproducibility. The automation platform included the following key components: An Agilent Bravo liquid handling workstation for accurate dispensing of reagents during titration and fixation. An Agilent VSpin centrifuge for gentle and programmable plate spinning to ensure even distribution and cell settling. A Thermo Scientific Cytomat automated incubator, specifically used to carry out incubation following PFA addition, ensuring consistent fixation conditions at controlled temperature. A temperature control pad is integrated with the deck layout to maintain localized cold zones. A multi-position plate hotel for temporary plate storage and routing across devices. Throughout the process, 96-well microplates were maintained at 4 °C on the temperature-controlled pad to preserve sample integrity during fixation. All steps were scheduled and coordinated through Cellario, ensuring seamless plate transfers and timing synchronization.

### TCRβ/αCD8 steric hindrance test

OTI CD8 T cells were isolated by negative selection (Miltenyi BIotec, 130-104-075) from OTI Thy1.1 spleens and mesenteric lymph nodes (Genentech in-house colony). Cells were first stained with live/dead aqua (ThermoFisher L34966). In one scenario, TCRb BV711 (BioLegend, 109243) was added first at 1:400 (that corresponds to 0.5 µg/mL or 0.05 µg per test) and incubated for 30 minutes at 4C, washed in MACS buffer, followed by anti-CD8 (clone CT-CD8a, ThermoFisher MA5-17594) at either 10 µg/mL or 50 µg/mL and incubated for an additional 30 minutes at 4C. In the second scenario, anti-CD8 was added first at either 10 µg/mL or 50 µg/mL and incubated for 30 minutes at 4C, washed, and then 1:400 of TCRb BV711 was added and incubated for an additional 30 minutes at 4C. Cells were washed twice in MACS buffer prior to flow cytometric analysis on a BD Symphony.

### Functional avidity test

OTI CD8 T cells were isolated by negative selection (Miltenyi BIotec, 130-104-075) from OTI Thy1.1 spleens and mesenteric lymph nodes (Genentech in-house colony). A portion of the spleen and lymph node cell suspensions were used as antigen presenting cells after T cell depletion. T cell depletion was done by first staining with anti-CD3 FITC (Tonbo Biosciences, 35-0032-U100), followed by removal using anti-FITC microbeads (Miltenyi Biotec, 130-048-701). Purified OTI CD8s and T-depleted APCs were plated at a 1:1 ratio with titrated purified peptides N4, T4, V4, UV peptide or VSV peptide. Peptide concentrations started from 10 µg/mL and serially diluted 10-fold across 12 wells. If CD8 blockade was included, anti-CD8 (clone CT-CD8a, ThermoFisher MA5-17594) was added at 50 µg/mL final. Cells were cultured for 4 hours with protein transport inhibitors (ThermoFisher, 00-4970-03). Cells were then harvested and stained with live/dead aqua (ThermoFisher L34966), TCRb BV711 (BioLegend, 109243), Thy1.1 PE (BioLegend 202524) and incubated ∼30 minutes at 4C. Cells were washed in MACS buffer then fixed in 1x fix/perm buffer (ThermoFisher, 00-5523-00 buffer kit), followed by 1x permeabilization and staining with IFNg APC (BioLegend, 505810) for 30 minutes at room temperature. Cells were washed in 1x perm buffer, followed by flow cytometric analysis on a BD Symphony.

### “TCR cycle” biokinetics model

The simple two-state model cannot account for the presence of the hump in the TCR-pMHC-CD8 binding kinetic curve. To explain this feature, we extend the basic reaction scheme to a slightly more complex model, as follows:

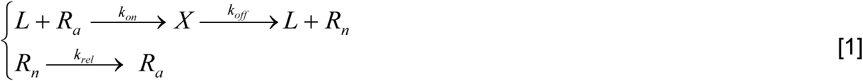

Here, *L* is the external ligand molecule; *R*_*a*_ and *R*_*n*_ are different types of free receptors, *X* is the ligand-receptor complex, and *k*_*j*_ are rate constants of corresponding reactions, described in detail in the legend of Figure 4.

The following differential equations describe the reaction scheme shown in [1]:

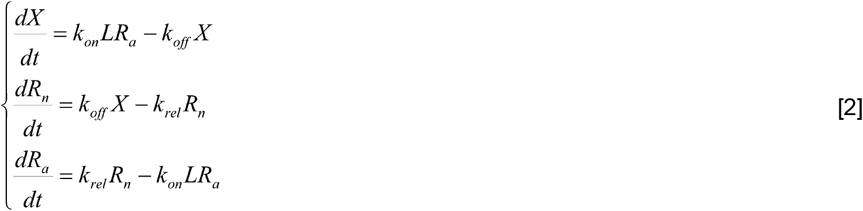

Equations [2] can be rewritten solely in terms of active receptors *R*_a_ and complexes *X*, by applying the conservation of the total number of receptors *R*_0_

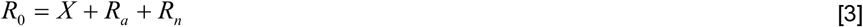

as follows:

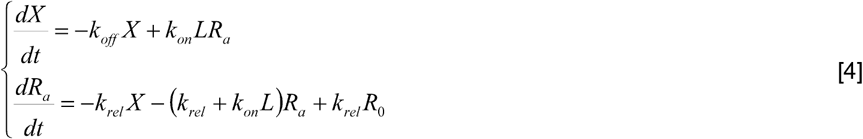

and solve it using the initial conditions for association kinetics:

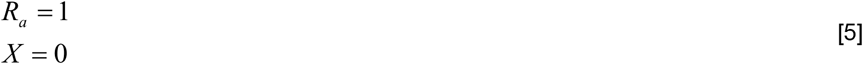

Or in matrix form:

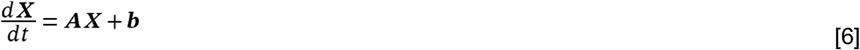

Where:

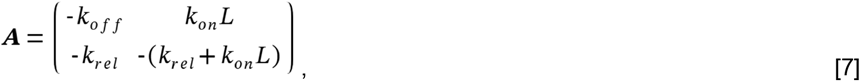

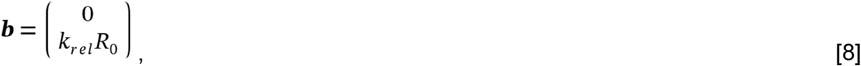

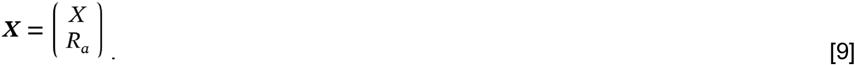

The matrix ***A*** has the following eigenvalues:

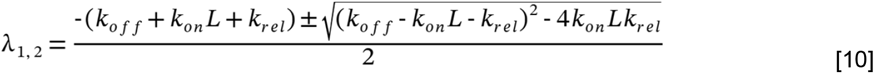

For example, for fitting with following optimal parameters:

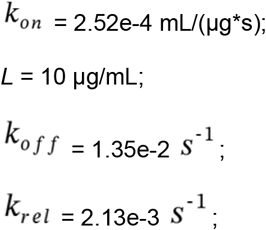

We have the matrix:

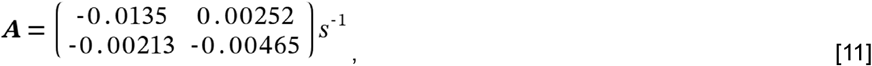

And the eigenvalues:

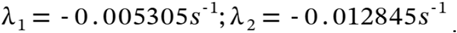

The intensity *Y(t)* of the MFI signal is then expressed in terms of the solution *X(t)* to the system of differential equations [4], as follows:

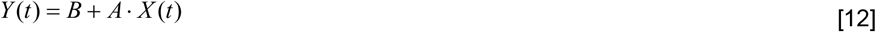

Here, *B* represents the background signal, and *A* = *C* × *R*_0_, where *C* is the instrument calibration constant (for the flow cytometer). The same notation is used for the concentrations of species as for the species themselves in the reaction scheme (e.g., *X*). In the fitting procedure, the fluorescence scaling factor *A* was intentionally held constant across conditions within each experimental batch. All OT-I–OVA measurements were acquired using identical flow cytometry settings within a batch (including PMT voltages and detector gains); accordingly, *A* represents a fixed experimental scaling factor rather than a condition-dependent biological variable. Allowing condition-specific values of *A* would violate the experimental design and introduce an unphysical degree of freedom capable of absorbing apparent kinetic differences unrelated to binding or competence cycling. For this reason, *A* was treated as a shared parameter by construction rather than optimized independently across conditions. In contrast, for datasets obtained from the public domain, *A* was fitted independently for each experimental condition, as the corresponding instrument settings were not reported and may have varied between experiments.

Reaction scheme [1] simplifies to the classic reversible receptor–ligand reaction (R + L ⇌ X) in the limit as 1/k_rel_ → 0. During fitting, we require the same value of *A* for the same batch of T cells, regardless of CD8 blocking or peptide type, since *A* = *C* × *R*_0_ represents the total number of receptors (*R*_0_) on the cell.

We developed a software package called *TCRkin* to fit binding kinetics data using the analytical solution of reaction scheme [1]:

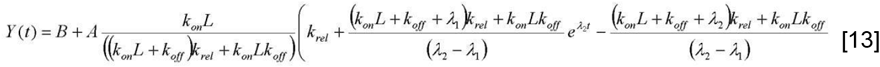

Where *λ*_*1*_ and *λ*_*2*_ are the eigenvalues:

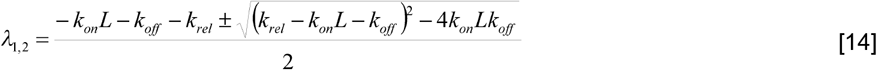

We emphasize that competence cycling is introduced here as a systems-level description of adaptive TCR behavior required to explain the observed kinetics. While intracellular phosphorylation-dependent regulation is a biologically plausible implementation, the model does not assume a unique molecular mechanism and is compatible with multiple biochemical realizations.

## Data availability

Source Data table in Leary et al [*Leary et al ., 2024*] is available through the journal web page: https://static-content.springer.com/esm/art%3A10.1038%2Fs41467-024-48198-0/Media_Objects/41467_2024_48198_MOESM4_ESM.xlsx

IMMREP_2022 benchmark data set [*Meysman et al ., 2024*] is available at https://github.com/viragbioinfo/IMMREP_2022_TCRSpecificity.

The OT1–OVA kinetic data newly generated as part of this study are publicly available at Zenodo: https://zenodo.org/records/17069783.

Digitized kinetic datasets reconstructed from previously published TCR–pMHC(HLA) studies analyzed in this manuscript (summarized in Figure 6e) are also publicly available at https://github.com/tabatsky/TCRkin/tree/main/digitized%20data. These datasets were generated using custom digitization tools developed by the authors, as described in the Code availability section.

## Code availability

The *TCRkin* source code, including the digitization algorithms used to reconstruct kinetic data from published figures, is available at https://github.com/tabatsky/TCRkin

## Acknowledgements

We acknowledge the use of OpenAI’s ChatGPT for assistance in refining the language and style of this manuscript. The model was utilized to improve clarity and coherence while ensuring that the original scientific content remained intact. All intellectual contributions and interpretations are the sole responsibility of the authors.

## Conflict of interests

MCT, SNP, MD, AS are employees of Genentech, Inc. IM is an employee of Medici Therapeutics. JC, MCT, MD, DO are inventors on a patent related to cellular assays and methods to assess TCR-pMHC interactions and kinetics.

## Contributions

Conceptualization and study design: MC, JD, JC, MCT, SNP, MD, RAS, AS, IM, AC, DO;

Data acquisition and curation: MC, JC, MCT, SNP, DO;

Methodology development: JC, MCT, SNP, MD, GS, AC, DO;

Data analysis and interpretation: MC, JD, JC, MCT, SNP, RAS, ET, AS, IM, AC, DO;

Manuscript drafting: MC, JD, JC, MCT, SNP, MD, RAS, ET, GS, AS, IM, AC, DO;

Critical review and editing: MC, JD, JC, MCT, SNP, MD, RAS, ET, GS, AS, IM, AC, DO;

Supervision and funding acquisition: IM, DO.

All authors have read and approved the article.

## Notes

### Summary of Updates

In the revised manuscript, we have substantially strengthened validation of the TCR cycle framework using several independently generated and previously published TCR-pMHC datasets that were not used in any way to develop, tune, or constrain the model (new Figure 6). We now include an explicit model comparison analysis demonstrating that the proposed competence-cycling mechanism is required by the data (new Figure S5). Specifically, fits of time-resolved binding kinetics show systematically improved goodness-of-fit when using the full TCR cycle model compared to a classic two-state reversible binding model. Model selection using Akaike Information Criterion consistently favors the TCR cycle model despite its additional parameter, particularly for kinetic curves exhibiting non-monotonic (hump-shaped) behavior. We further report parameter correlation and identifiability analyses, clarifying how kon, koff, and krel are constrained by the experimental design. Together, these results demonstrate that competence cycling is not a speculative addition but a quantitatively justified mechanistic requirement. We have substantially expanded the Supplementary Information to provide a detailed, experimentally grounded protocol for the proposed library-on-library kinetic data generation framework. The revised text now explicitly describes (i) the barcoding strategy, including how TCR identity, ligand identity, and time-point information can be uniquely encoded, (ii) the sequencing and/or flow-cytometry-based readout workflow, (iii) expected experimental throughput, and (iv) feasibility and approximate cost considerations. Together, these additions are intended to allow readers to clearly evaluate the practical implementation of the approach.

