## Supplementary Materials for "Tricked by Edge Cases: Can Current Approaches Lead to Accurate Prediction of T-Cell Specificity with Machine Learning?"

### This PDF file includes:

Supplementary Text
Figures. S1 to S6
Tables S1 to S2
Captions for Data S1 to S2

### Other Supplementary Materials for this manuscript include the following:

Data S1 to S2

### Supplementary text

#### Condition for hump formation

The condition for hump existence is a zero first derivative and a negative second derivative of the signal.

First derivative equals zero:

$$\frac{d}{dt}Y(t)=0=A\frac{k_{rel}k_{on}L}{\left(\left(k_{on}L+k_{off}\right)k_{rel}+k_{on}Lk_{off}\right)}\left(\lambda_2\frac{\left(k_{on}L+k_{off}+\lambda_1\right)k_{rel}+k_{on}Lk_{off}}{\left(\lambda_2-\lambda_1\right)k_{rel}}e^{\lambda_2 t}-\lambda_1\frac{\left(k_{on}L+k_{off}+\lambda_2\right)k_{rel}+k_{on}Lk_{off}}{\left(\lambda_2-\lambda_1\right)k_{rel}}e^{\lambda_1 t}\right) \quad [1]$$

$$\lambda_2\frac{\left(k_{on}L+k_{off}+\lambda_1\right)k_{rel}+k_{on}Lk_{off}}{\left(\lambda_2-\lambda_1\right)k_{rel}}e^{\lambda_2 t}=\lambda_1\frac{\left(k_{on}L+k_{off}+\lambda_2\right)k_{rel}+k_{on}Lk_{off}}{\left(\lambda_2-\lambda_1\right)k_{rel}}e^{\lambda_1 t} \quad [2]$$

$$e^{(\lambda_2-\lambda_1)t}=\frac{\lambda_1}{\lambda_2}\cdot\frac{\left(k_{on}L+k_{off}+\lambda_2\right)k_{rel}+k_{on}Lk_{off}}{\left(k_{on}L+k_{off}+\lambda_1\right)k_{rel}+k_{on}Lk_{off}} \quad [3]$$

Condition of negative second derivative:

$$\frac{d^2}{dt^2}Y(t)=A\frac{k_{rel}k_{on}L}{\left(\left(k_{on}L+k_{off}\right)k_{rel}+k_{on}Lk_{off}\right)}\left(\lambda_2^2\frac{\left(k_{on}L+k_{off}+\lambda_1\right)k_{rel}+k_{on}Lk_{off}}{\left(\lambda_2-\lambda_1\right)k_{rel}}e^{\lambda_2 t}-\lambda_1^2\frac{\left(k_{on}L+k_{off}+\lambda_2\right)k_{rel}+k_{on}Lk_{off}}{\left(\lambda_2-\lambda_1\right)k_{rel}}e^{\lambda_1 t}\right)<0 \quad [4]$$

$$\lambda_2^2\frac{\left(k_{on}L+k_{off}+\lambda_1\right)k_{rel}+k_{on}Lk_{off}}{\left(\lambda_2-\lambda_1\right)k_{rel}}e^{\lambda_2 t}-\lambda_1^2\frac{\left(k_{on}L+k_{off}+\lambda_2\right)k_{rel}+k_{on}Lk_{off}}{\left(\lambda_2-\lambda_1\right)k_{rel}}e^{\lambda_1 t}<0 \quad [5]$$

1) For real (non-oscillatory) solutions $\lambda_1>\lambda_2$, $\lambda_1<0$, $\lambda_2<0$ :

$$\lambda_2\lambda_1 \frac{(k_{on}L + k_{off} + \lambda_2)k_{rel} + k_{on}Lk_{off}}{(\lambda_2 - \lambda_1)k_{rel}} e^{\lambda_1 t} - \lambda_1^2 \frac{(k_{on}L + k_{off} + \lambda_2)k_{rel} + k_{on}Lk_{off}}{(\lambda_2 - \lambda_1)k_{rel}} e^{\lambda_1 t} < 0 \quad [6]$$

$$\lambda_2(k_{on}L + k_{off} + \lambda_2)k_{rel} + \lambda_2 k_{on}Lk_{off} < \lambda_1(k_{on}L + k_{off} + \lambda_2)k_{rel} + \lambda_1 k_{on}Lk_{off} \quad [7]$$

$$0 < (\lambda_1 - \lambda_2)(k_{on}L + k_{off} + \lambda_2)k_{rel} + (\lambda_1 - \lambda_2)k_{on}Lk_{off} \quad [8]$$

$$(k_{on}L + k_{off} + \lambda_2)k_{rel} + k_{on}Lk_{off} > 0 \quad [9]$$

$$\lambda_2 > -\frac{k_{on}Lk_{off}}{k_{rel}} - k_{on}L - k_{off} \quad [10]$$

$$\frac{-k_{on}L - k_{off} - k_{rel} - \sqrt{(k_{rel} - k_{on}L - k_{off})^2 - 4k_{on}Lk_{off}}}{2} > -\frac{k_{on}Lk_{off}}{k_{rel}} - k_{on}L - k_{off} \quad [11]$$

$$-\sqrt{(k_{rel} - k_{on}L - k_{off})^2 - 4k_{on}Lk_{off}} > -\frac{2k_{on}Lk_{off}}{k_{rel}} - k_{on}L - k_{off} + k_{rel} \quad [12]$$

$$\begin{cases} \sqrt{(k_{on}L + k_{off} - k_{rel})^2 - 4k_{on}Lk_{off}} < \dfrac{2k_{on}Lk_{off}}{k_{rel}} + k_{on}L + k_{off} - k_{rel} \\ (k_{on}L + k_{off} - k_{rel})^2 - 4k_{on}Lk_{off} > 0 \end{cases} \quad [13]$$

This ultimately leads to the following condition:

$$k_{rel} < k_{on}L + k_{off} - \sqrt{4k_{on}Lk_{off}} \quad [14]$$

Thus, the condition for hump formation is given by equation [14], which corresponds to very slow receptor relaxation following complex dissociation.

2) Complex (oscillatory) solutions $\lambda_1$ and $\lambda_2$ consistently exhibit a hump:

$$(k_{rel} - k_{on}L - k_{off})^2 - 4k_{on}Lk_{off} < 0 \quad [15]$$

This leads to the following condition:

$$k_{on}L + k_{off} - \sqrt{4k_{on}Lk_{off}} < k_{rel} < k_{on}L + k_{off} + \sqrt{4k_{on}Lk_{off}} \quad [16]$$

Thus, taking both cases into account, the hump condition is given by:

$$k_{rel} < k_{on}L + k_{off} + \sqrt{4k_{on}Lk_{off}} \quad [17]$$

#### Multiplexed library-on-library measurement of TCR–pHLA(MHC) binding kinetics

This section outlines a proposed experimental strategy and does not represent data generated or analyzed in the present study.

##### Overview

To enable scalable inference of TCR–pHLA association and dissociation kinetics, we outline a multiplexed library-on-library framework, enabling tens to hundreds of receptors to be interrogated against hundreds of ligands in a single experiment. This approach integrates transient TCR expression via mRNA electroporation, DNA-barcoded monomeric pHLAs, time-point–specific barcoding, and downstream flow-cytometric and sequencing-based readouts to reconstruct binding-kinetics curves and estimate the kinetic parameters *kon*, *koff, krel*.

##### Generation of TCR-expressing primary $CD8^+$ T cells

Primary human $CD8^+$ T cells can be isolated from peripheral blood mononuclear cells by negative selection. Individual TCRαβ constructs are synthesized as in vitro–transcribed mRNA and introduced into $CD8^+$ T cells by electroporation. Each electroporation yields a homogeneous population of approximately $10^5$–$10^6$ viable T cells expressing a single, defined TCR. To preserve TCR identity after pooling, each TCR population is labeled with a unique cell-intrinsic barcode, implemented either as (i) a DNA hashtag oligonucleotide or (ii) a short genetic barcode co-electroporated alongside the TCR mRNA.

##### Pooling of TCR libraries

Following recovery and verification of surface TCR expression, all TCR-expressing cell populations can be pooled into a single suspension (Figure S6a). Pooling prior to ligand exposure ensured that all TCR–pHLA interactions were measured under identical staining, temperature, and handling conditions, minimizing batch effects and enabling direct comparison of kinetic parameters across receptors.

#### DNA-barcoded monomeric pHLA library

Recombinant monomeric pHLAs can be generated with site-specific conjugation to DNA oligonucleotide barcodes, each uniquely identifying a peptide–HLA species. All pHLAs are labeled with a common fluorophore to enable quantitative detection of binding by flow cytometry. The attached DNA barcodes enable multiplexed decoding of ligand identity during downstream sequencing.

#### Time-resolved binding, activation, and barcoding

For measurements focused exclusively on TCR–pHLA binding kinetics, pooled TCR-expressing cells can be incubated with a pooled, DNA-barcoded pHLA library under controlled temperature and concentration conditions (Figure S6b). Binding is initiated at time zero and quenched at defined time points by the addition of 4% paraformaldehyde. At each time point, cells are fixed and labeled with a unique, time-point–specific barcode, enabling post hoc reconstruction of association and dissociation kinetics.

In contrast, for experiments aimed at simultaneously measuring TCR triggering or early T cell activation, pooled TCR-expressing cells must be incubated with individual pHLA ligands in separate reactions (Figure S6c). This design is required to unambiguously attribute downstream signaling or activation readouts to a specific pHLA, which would not be possible in a fully pooled ligand condition.

#### Flow cytometry and sequencing readout

Cells can be analyzed either by flow cytometry to quantify fluorescence intensity corresponding to pHLA binding at each time point, or by sequencing-based readout following pooling of samples. In the sequencing-based approach, TCR identity barcodes, pHLA DNA barcodes, and time-point–specific barcodes are recovered from the same cells. Together, these measurements generate a dataset linking TCR sequence, ligand identity, binding signal, and time.

While flow cytometry alone can be used to measure binding kinetics for small numbers of predefined TCR–pHLA pairs, scalable deconvolution of TCR identity, ligand identity, and time point requires a sequencing-based barcoding strategy. Flow cytometry provides quantitative binding readout, whereas DNA barcodes enable multiplexed decoding of interaction identity.

In the sequencing-based implementation, the amount of pHLA bound at each time point is inferred from normalized counts of pHLA-conjugated DNA barcodes recovered from cells, analogous to CITE-seq–based quantification of surface protein abundance.

#### Kinetic modeling and parameter inference

For each TCR–pHLA pair, the binding signal as a function of time could be reconstructed and fitted to the appropriate biophysical kinetic model.

#### Feasibility, throughput, and cost considerations

The proposed multiplexed library-on-library kinetic framework is experimentally tractable using established immunology technologies. Transient mRNA electroporation of primary human $CD8^+$ T cells reliably yields $10^5$–$10^6$ viable cells per TCR construct within 1–2 days, allowing parallel generation of dozens of distinct TCR populations without viral transduction or clonal expansion.

DNA-barcoded monomeric pHLAs can be produced or sourced at scale using existing conjugation chemistries, and hundreds of ligands can be pooled and assayed simultaneously in a single tube or 96-well format.

For example (Table S2), a single experiment could include 50 unique TCRs × 200 distinct pHLAs, representing 10,000 TCR–pHLA interactions measured across multiple time points. Time-point barcoding eliminates the need for separate wells per kinetic condition, reducing reagent usage and experimental complexity. In practice, the limiting factor is sequencing depth rather than wet-lab throughput. Estimated per-experiment costs are dominated by oligonucleotide barcodes and sequencing, which scale sublinearly with library size, making this approach substantially more cost-effective than measuring each TCR–pHLA pair individually. Together, these features render the method feasible, scalable, and well suited for generating the large kinetic datasets required for data-driven or machine-learning–based modeling of TCR–pHLA interactions.

The resulting large-scale, time-resolved kinetic measurements enable supervised and physics-informed machine-learning models that map TCR and pHLA sequences directly to the rate constants of the TCR cycle model, including the association *kon*, dissociation *koff*, and *krel* rates. In particular, transformer-based sequence encoders and physics-informed neural networks can be trained in a multi-task setting to predict these parameters while enforcing the underlying biophysical kinetics, providing a principled framework for learning sequence–kinetics relationships beyond equilibrium affinity.

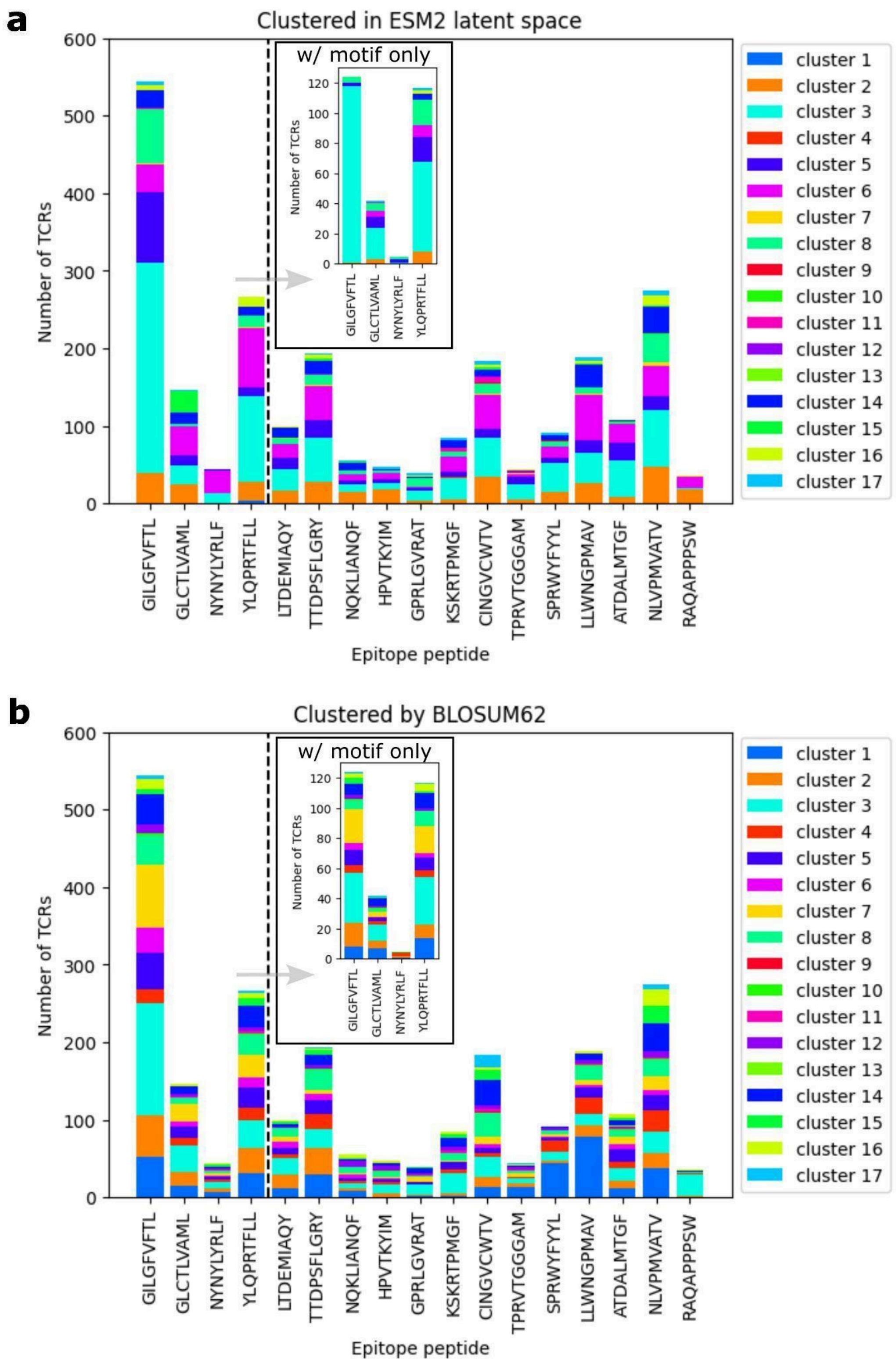
a
Clustered in ESM2 latent space
w/ motif only
Number of TCRs
Epitope peptide
cluster 1
cluster 2
cluster 3
cluster 4
cluster 5
cluster 6
cluster 7
cluster 8
cluster 9
cluster 10
cluster 11
cluster 12
cluster 13
cluster 14
cluster 15
cluster 16
cluster 17
GILGFVFTL
GLCTLVAML
NYNYLYRLF
YLQPRTFLL
LTDEMIAQY
TTDPSFLGRY
NQKLIANQF
HPVTKYIM
GPRLGVRAT
KSKRTPMGF
CINGVCWTV
TPRVTGGGAM
SPRWYFYYL
LLWNGPMAV
ATDALMTGF
NLVPMVATV
RAQAPPPSW
b
Clustered by BLOSUM62
w/ motif only
Number of TCRs
Epitope peptide

**Figure. S1. Clustering of TCRs in protein language model or sequence space does not lead to grouping by their epitope specificity.** Distribution of TCRs from 17 peptide-specific data sets from IMMREP_2022 benchmark in 17 clusters acquired with hierarchical clustering based on TCR$\beta$ distance calculated using mean ESM2 (3B variant - 2560 dimensions) latent representations of TCR CDR3$\beta$ sequences (**a**) and BLOSUM62 global alignment score among TCR CDR3$\beta$ sequences (**b**). First four peptides separated by the dashed line contain a significant CDR3$\beta$ binding motif. Distribution of motif-containing-only data points in the 17 clusters is shown in the inset plots (motifs detected using sequence logos).

**a**

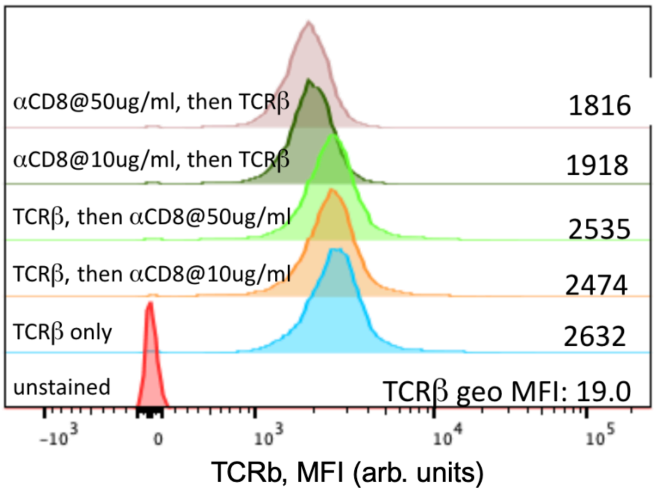
αCD8@50ug/ml, then TCRβ
1816
αCD8@10ug/ml, then TCRβ
1918
TCRβ, then αCD8@50ug/ml
2535
TCRβ, then αCD8@10ug/ml
2474
TCRβ only
2632
unstained
TCRβ geo MFI: 19.0
-10^3
0
10^3
10^4
10^5
TCRb, MFI (arb. units)

**b**

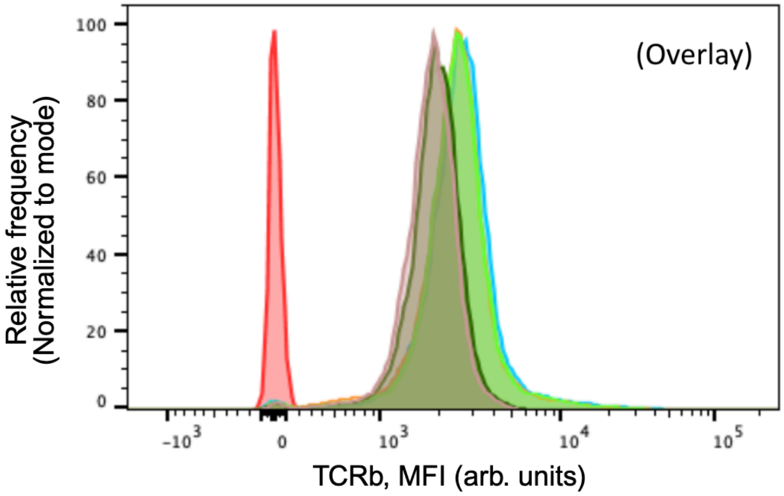
100
80
60
40
20
0
Relative frequency
(Normalized to mode)
(Overlay)
-10^3
0
10^3
10^4
10^5
TCRb, MFI (arb. units)

**Figure. S2. TCRβ/αCD8 steric hindrance test on purified OTI CD8 T cells–prebound αCD8 partially blocks αTCRβ binding.**

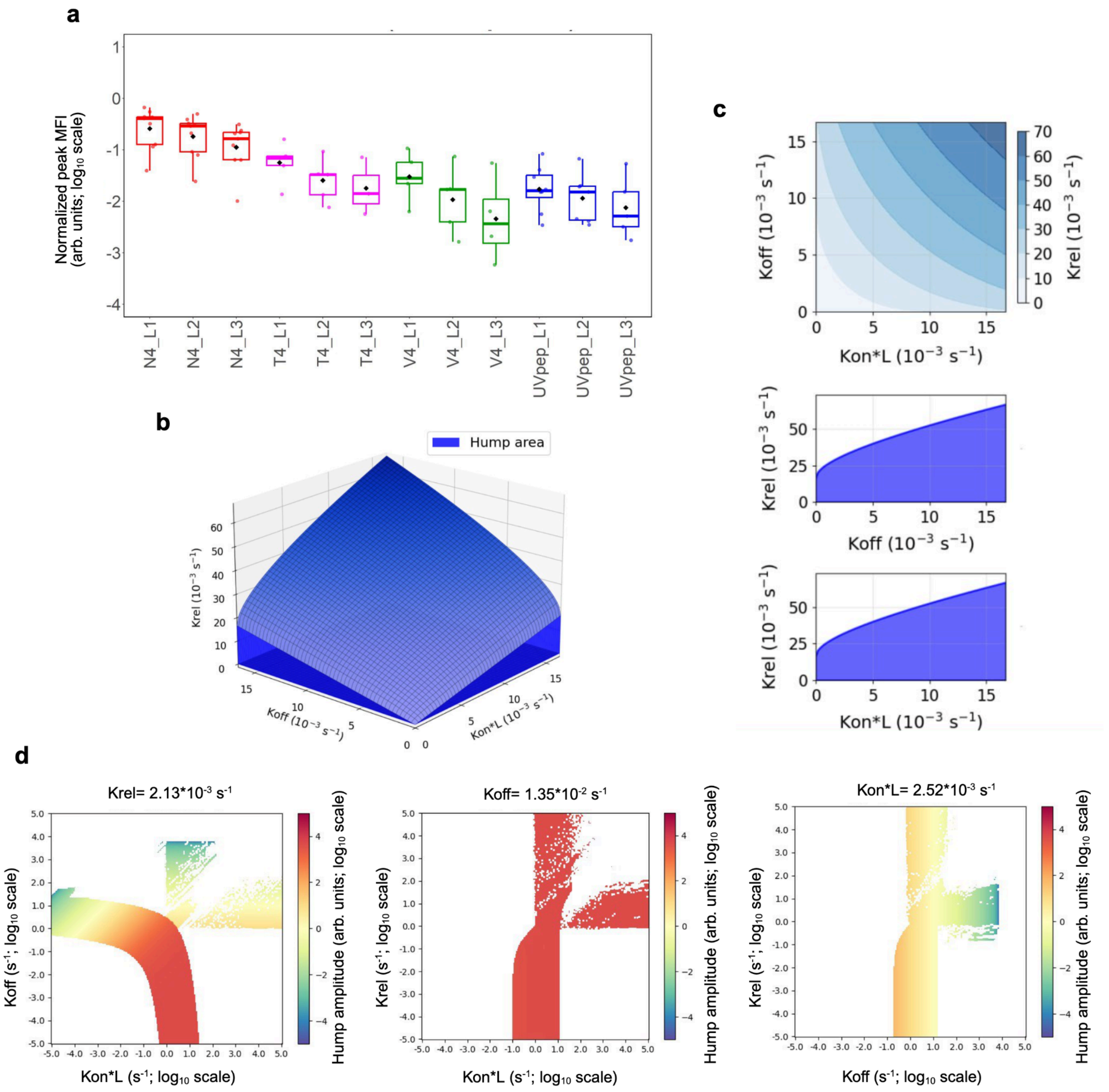

a
Normalized peak MFI (arb. units; log10 scale)
N4_L1
N4_L2
N4_L3
T4_L1
T4_L2
T4_L3
V4_L1
V4_L2
V4_L3
UVpep_L1
UVpep_L2
UVpep_L3
b
Hump area
Krel (10^-3 s^-1)
Koff (10^-3 s^-1)
Kon*L (10^-3 s^-1)
c
Koff (10^-3 s^-1)
Kon*L (10^-3 s^-1)
Krel (10^-3 s^-1)
Krel (10^-3 s^-1)
Koff (10^-3 s^-1)
Krel (10^-3 s^-1)
Kon*L (10^-3 s^-1)
d
Krel= 2.13*10^-3 s^-1
Koff (s^-1; log10 scale)
Kon*L (s^-1; log10 scale)
Hump amplitude (arb. units; log10 scale)
Koff= 1.35*10^-2 s^-1
Krel (s^-1; log10 scale)
Kon*L (s^-1; log10 scale)
Hump amplitude (arb. units; log10 scale)
Kon*L= 2.52*10^-3 s^-1
Krel (s^-1; log10 scale)
Koff (s^-1; log10 scale)
Hump amplitude (arb. units; log10 scale)

**Figure. S3. Peak binding features and kinetic parameter regimes associated with hump formation in TCR–pMHC binding kinetics. a**. Peak MFI values observed during TCR–pMHC binding for three pMHC concentrations (L1, L2, and L3 corresponding to 10, 5, and 2 μg/mL, respectively). Data are shown on a $\log_{10}$ scale. To enable comparison across experimental days, MFI values were normalized using (peak − B) / A, where *A* and *B* correspond to flow cytometer scaling parameters. Error bars represent the standard deviation of values across independent experiments. **b**. Region of kinetic parameter space (*kon*, *koff*, *krel*) that gives rise to a non-monotonic "hump" in the binding kinetics curve, highlighted in blue. The boundary conditions for hump formation are derived analytically and described in the Supplementary Materials. **c**. Two-dimensional projections of the three-dimensional parameter space shown in panel b, illustrating pairwise relationships between *kon*, *koff*, and *krel* that support hump formation. **d**. Two-dimensional phase diagrams generated using kinetic parameters inferred from a representative OT1–OVA–N4 binding experiment, illustrating the region of parameter space in which the hump-forming regime is present; hump amplitude is shown using color coding.

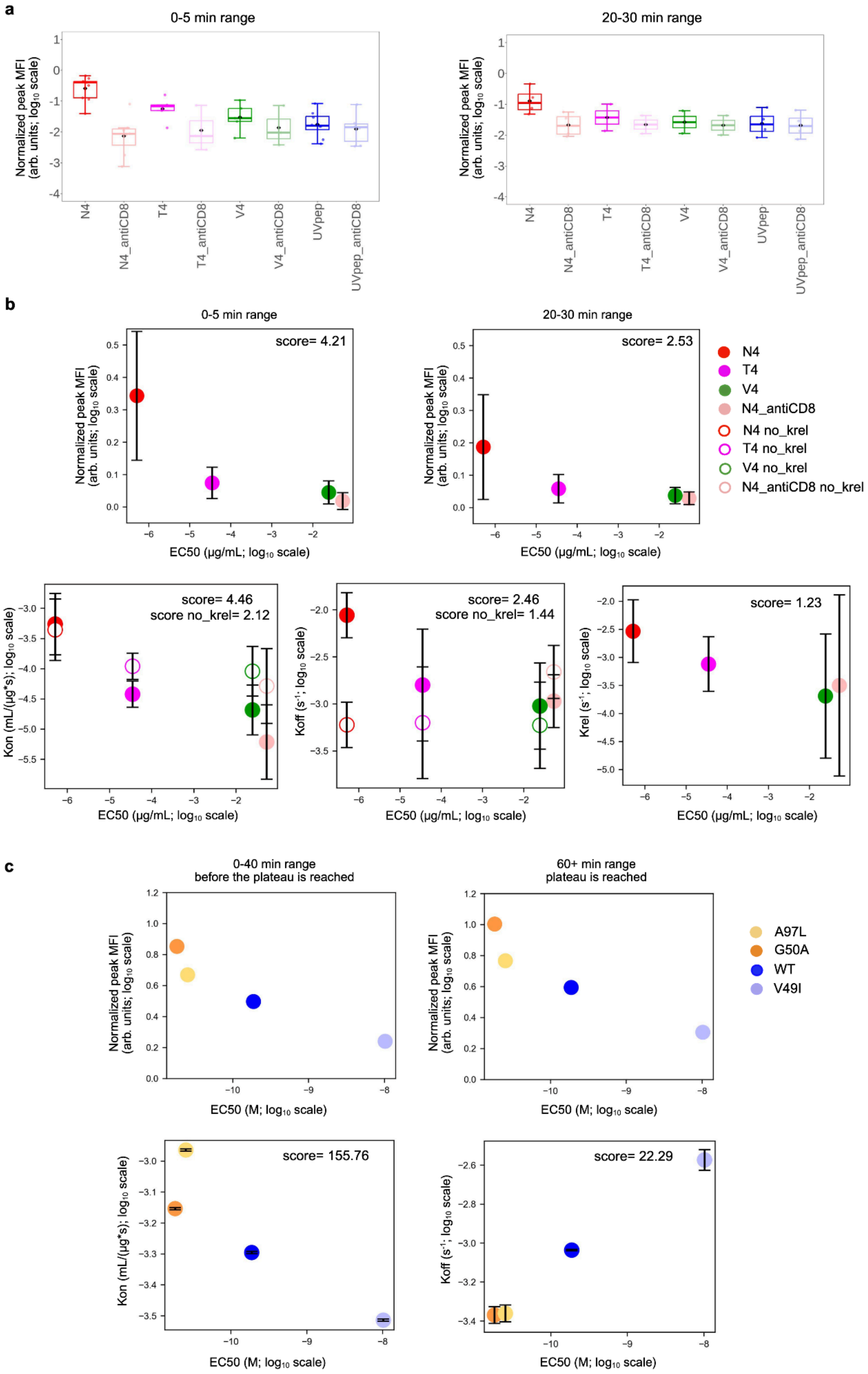
a
0-5 min range
20-30 min range
Normalized peak MFI (arb. units; log10 scale)
N4
N4_antiCD8
T4
T4_antiCD8
V4
V4_antiCD8
UVpep
UVpep_antiCD8
b
0-5 min range
20-30 min range
score= 4.21
score= 2.53
EC50 (μg/mL; log10 scale)
N4
T4
V4
N4_antiCD8
N4 no_krel
T4 no_krel
V4 no_krel
N4_antiCD8 no_krel
score= 4.46
score no_krel= 2.12
score= 2.46
score no_krel= 1.44
score= 1.23
Kon (mL/(μg*s); log10 scale)
Koff (s-1; log10 scale)
Krel (s-1; log10 scale)
c
0-40 min range
before the plateau is reached
60+ min range
plateau is reached
EC50 (M; log10 scale)
A97L
G50A
WT
V49I
score= 155.76
score= 22.29

**Figure. S4. Early binding features better predict TCR functional avidity. a**. Comparison of peak binding signals (mean fluorescence intensity, MFI) measured at early (0–5 min) versus late (20–30 min) time windows for OVA and mutant pMHC ligands. For each peptide, peak values were extracted from the measured TCR–pMHC association kinetics and summarized across replicates. Early-time peaks show greater separation between OVA and mutant ligands (**b**) than late-time peaks. Discriminability was quantified using a normalized effect-size score defined as *(max(mean) − min(mean)) / average SD*, with uncertainty estimated across replicates. Kinetic parameters inferred using the competence cycle model (including krel) show superior discriminability compared to a simple reversible binding model lacking competence cycling (no_krel). Correlations between fitted kinetic parameters (kon,koff,krel) and functional avidity ($EC_{50}$), operationally defined as the $EC_{50}$ from cytokine production assays, are shown. Error bars represent the standard deviation of values across independent experiments. **c.** Correlation of peak binding values measured at early (0–40 min) versus late (≥60 min) time windows for the NY–ESO-$1_{157–165}$ peptide presented by HLA-A*0201 and variant TCRs, as reported in [*Schmid et al., 2010*], with functional avidity ($EC_{50}$ for target cell lysis) and inferred kinetic parameters. *koff* values were independently obtained by fitting the empirically measured dissociation curves from Figure 1c in [*Schmid et al., 2010*] using our model, which, in the dissociation regime, reduces to the standard single-exponential decay model. These *koff* values were then used as fixed input parameters to fit the association kinetics curves within the TCR cycle model. Error bars represent the standard errors of the fitted parameters, derived from the Hessian matrix of the fit.

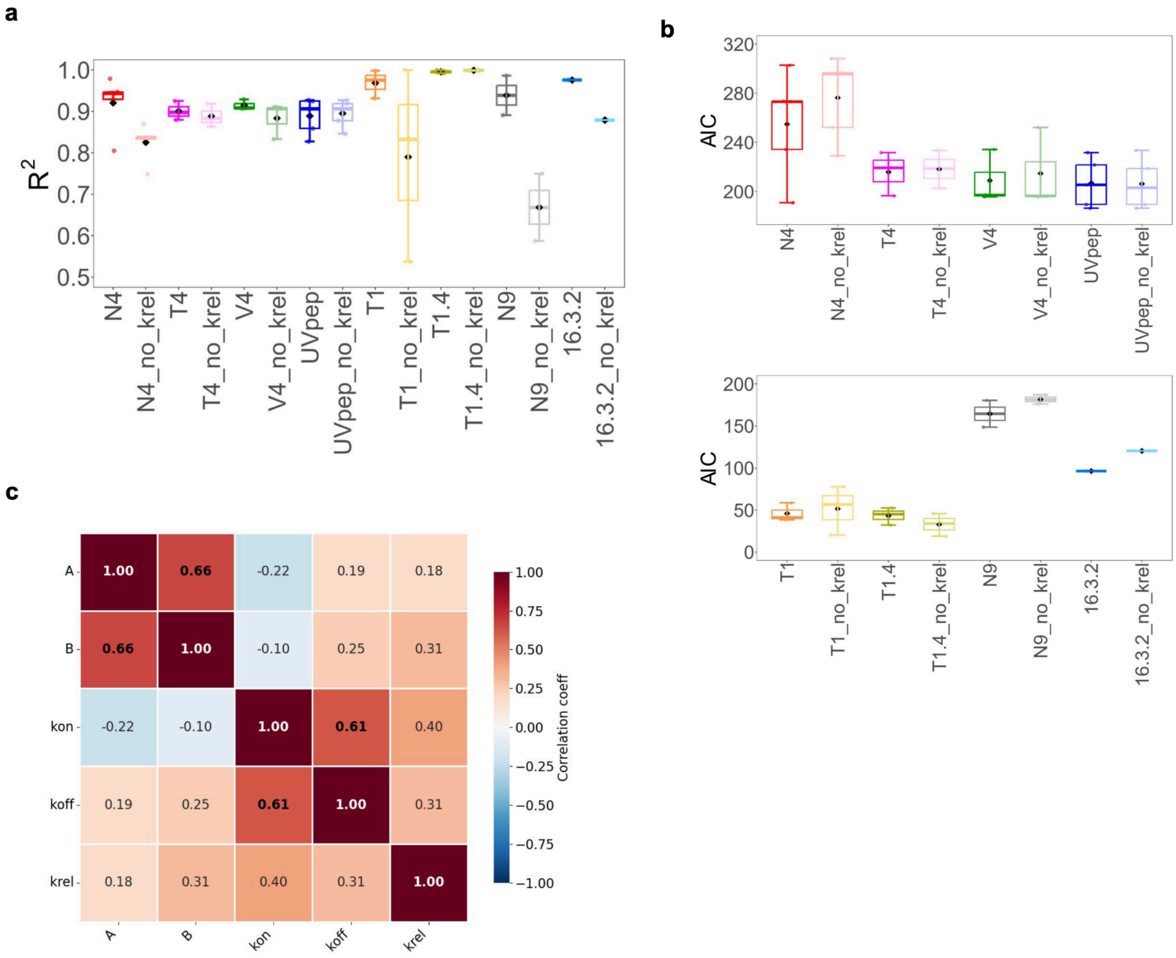
a
b
c
R2
1.0
0.9
0.8
0.7
0.6
0.5
N4
N4_no_krel
T4
T4_no_krel
V4
V4_no_krel
UVpep
UVpep_no_krel
T1
T1_no_krel
T1.4
T1.4_no_krel
N9
N9_no_krel
16.3.2
16.3.2_no_krel
AIC
320
280
240
200
150
100
50
0
A
B
kon
koff
krel
1.00
0.66
-0.22
0.19
0.18
0.66
1.00
-0.10
0.25
0.31
-0.22
-0.10
1.00
0.61
0.40
0.19
0.25
0.61
1.00
0.31
0.18
0.31
0.40
0.31
1.00
1.00
0.75
0.50
0.25
0.00
−0.25
−0.50
−0.75
−1.00
Correlation coeff

**Figure. S5. Model comparison supports the necessity of competence cycling in TCR–pMHC kinetics. a**. Goodness-of-fit ($R^2$) for time-resolved TCR–pMHC binding data fitted with the full TCR cycle model incorporating ligand release from the competent state (*krel*) versus a classic two-state reversible binding model lacking competence cycling ("no_*krel*"). Across ligands and conditions, inclusion of *krel* systematically improves fit quality. **b**. Model comparison using Akaike Information Criterion (AIC) demonstrates a consistent preference for the TCR cycle model over the reversible two-state model, despite the additional parameter, indicating that competence cycling is warranted by the data, particularly for kinetic curves that exhibit a pronounced hump. **c**. Correlation matrix of fitted parameters for the TCR cycle model, illustrating parameter dependencies and practical identifiability under the experimental time windows and ligand concentration ranges used in OT1-OVA system. Error bars represent the standard deviation of values across independent experiments.

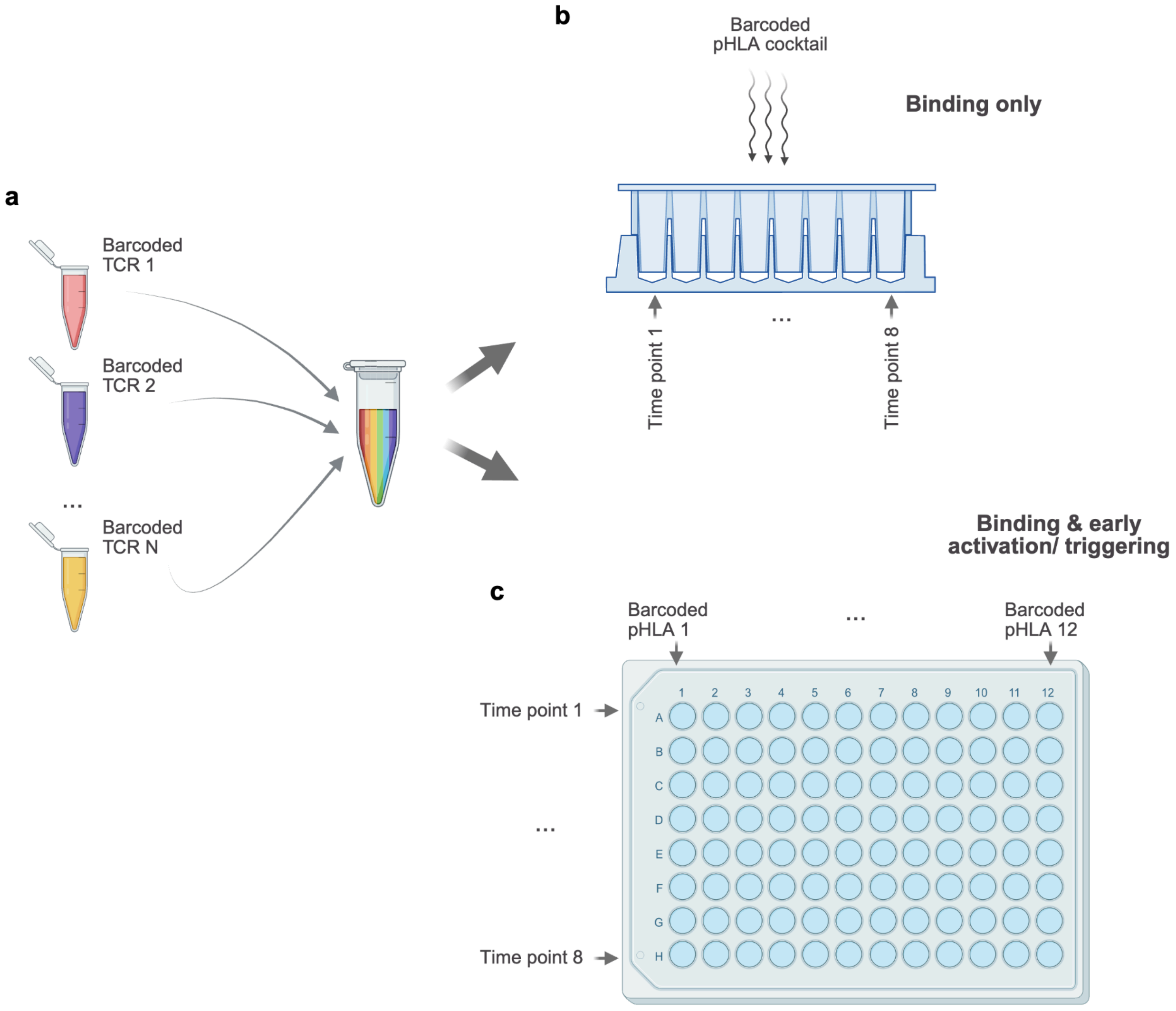

a
Barcoded TCR 1
Barcoded TCR 2
...
Barcoded TCR N
b
Barcoded pHLA cocktail
Binding only
Time point 1
...
Time point 8
Binding & early activation/ triggering
c
Barcoded pHLA 1
...
Barcoded pHLA 12
Time point 1
...
Time point 8
1 2 3 4 5 6 7 8 9 10 11 12
A B C D E F G H

**Figure. S6. Multiplexed experimental designs for scalable TCR–pHLA(MHC) kinetic and functional measurements. a.** Generation and pooling of TCR-expressing primary human CD8$^+$ T cells. Individual TCRαβ constructs are transiently expressed via mRNA electroporation, and each TCR population is labeled with a unique cell-intrinsic barcode prior to pooling. **b.** Fully pooled library-on-library configuration for TCR–pHLA binding kinetics. Barcoded TCR-expressing cells are incubated with a pooled library of DNA-barcoded monomeric pHLA ligands under identical biochemical conditions. Binding is initiated at time zero and quenched at defined time points, which are encoded using time-point–specific barcodes, enabling simultaneous reconstruction of association and dissociation kinetics across many TCR–pHLA pairs. **c.** Partially pooled configuration for measuring binding together with early T cell activation. Barcoded TCR-expressing cells are pooled as in (**a**) but incubated with individual pHLA ligands in separate reactions, allowing unambiguous attribution of downstream signaling or activation readouts to a specific pHLA. Figure elements created with BioRender.com

| Type of an observation | Assay type | Conclusion | Reference |
|---|---|---|---|
| Binding or activation following a multimer stain or incubation with peptides is not an indicator of whether a T cell is specific to an endogenously presented epitope | Peptide-based MIRA | Multimer staining or incubation with peptides is not an indicator of T cell specificity against endogenously presented epitopes | Nolan et al [*Nolan et al., 2020*] |
| Up to 72% of TCR clones against C virus NS3 variant peptides as well as the HY self-antigen were tetramer positive yet did not produce an antigen specific response | Tetramer staining and functional response by CD107a | pMHC binding does not uniformly predict T cell activity | Sibener et al Yu et al [*Sibener et al., 2018; Yu et al., 2015*] |
| pMHC class I tetramers and dextramers can fail to detect relevant functional T cell clonotypes and underestimate antigen-reactive T cell populations | Multimer staining and functional assays including cytokine release and cytotoxic assays<br><br>Dodecamers vs dextramers vs tetramers | Many previous attempts to enumerate antigen specific T cell populations by pMHC multimer staining will have underestimated the size of these populations | Rius et al Tungatt et al Huang et al [*Rius et al., 2018; Tungatt et al., 2015; Huang et al., 2016*] |

**Table S1. Representative studies underscore how multimer-binding assays significantly bias our perception of TCR specificity.**

| Parameter | Value/Example | Notes |
|---|---|---|
| Number of TCRs per experiment | 50 (using 96 well EP plate) | Primary human $CD8^+$ T cells electroporated with individual TCR mRNAs |
| Number of pHLAs per experiment | 200 | DNA-barcoded monomeric pHLAs, pooled for multiplexed staining |
| Total TCR–pHLA interactions | 10,000 | 50 TCRs × 200 pHLAs |
| Cells per TCR | $5–10 \times 10^5$ | Sufficient to maintain statistical power and FACS resolution |
| Total cells per experiment | $5–10 \times 10^6$ (adjust if needed) | Easily achievable with standard PBMC isolation and expansion |
| Time points per kinetic curve | 5–8 | Association measurements; barcoded for sequencing |
| Sequencing reads required | ~50–100k per TCR–pHLA–time point | Ensures robust reconstruction of binding curves |
| Estimated cost | $10–25k per experiment | Includes TCR mRNA synthesis, pHLA production, barcoding, and sequencing; scales sublinearly with library size |
| Experimental duration | Up to 2 weeks: ~10 days for TCR generation, and 2-4 days for seq prep | Includes T cell recovery, electroporation, staining, fixation, and sequencing preparation |

**Table S2. Representative experimental scale, throughput, and resource requirements for the multiplexed TCR–pHLA kinetic measurement framework.** The table summarizes typical parameter values for a single multiplexed experiment, including the number of TCRs and pHLA ligands, total interaction space, cell numbers, time-point resolution, cost, and duration. Values are provided as representative examples and may be adjusted depending on experimental design, library composition, and sequencing depth requirements.

**Data S1**. **(separate file)**
Parameter values with corresponding multiplicative errors (e.g., */ 1.01e+00 indicates that the value may vary by a factor of 1.01, or approximately ±1%, from the best-fit estimate) obtained from fitting the TCR cycle model to experimental data collected on different experimental days (see 'fileName' column).

**Data S2**. **(separate file)**
Average MFI values (arbitrary units) of pTCR signal measured in three independent experiments.
